# Distinct metabolic states of a cell guide alternate fates of mutational buffering through altered proteostasis

**DOI:** 10.1101/540039

**Authors:** Kanika Verma, Kanika Saxena, Rajashekar Donaka, Aseem Chaphalkar, Manish Kumar Rai, Anurag Shukla, Zainab Zaidi, Rohan Dandage, Dhanasekaran Shanmugam, Kausik Chakraborty

**Author notes:** Equally contributing first authors (Kanika Verma, Kanika Saxena). Institute of Biomedicine, University of Gothenburg, Medicinaregatan 7A, Gothenburg, Sweden (Kanika Saxena).

## Abstract

Changes in metabolism can alter the cellular milieu; can this also change intracellular proteostasis? Since proteostasis can modulate mutational buffering, if change in metabolism has the ability to change proteostasis, arguably, it should also alter mutational buffering. Building on this, we find that altered cellular metabolic states in *E. coli* buffer distinct mutations. Buffered-mutants had folding problems *in vivo* and were differently chaperoned in different metabolic states. Notably, this assistance was dependent upon the metabolites and not on the increase in canonical chaperone machineries. Additionally, we were able to reconstitute the folding assistance afforded by metabolites *in vitro* and propose that changes in metabolite concentrations have the potential to alter proteostasis. Collectively, we unravel that the metabolite pools are bona fide members of proteostasis and aid in mutational buffering. Given the plasticity in cellular metabolism, we posit that metabolic alterations may play an important role in the positive or negative regulation of proteostasis.

---

Metabolic rewiring is a common response among different organisms to their surrounding environment^1^. Different cell types differ in their preferred mode of metabolism in order to harness energy and generate its required set of metabolites^2–5^. Interestingly, many of the cellular metabolites are known to change protein folding pathways by modifying protein stability and folding kinetics *in vitro* at high concentrations. Some of the special ones like polyphosphates^6,7^ and ATP^8,9^ are known to affect cellular protein aggregation during certain stresses. It is important to know if the change in metabolite composition of cellular milieu can alter intracellular protein folding capacity.

If metabolites and metabolism can affect proteostasis it may have two fundamental implications. 1) Metabolism-dependent change in proteostasis may aid evolution of the whole proteome when there is a change in an organism’s niche or surrounding climate that alters its internal environment, or when an organism undergoes a large change in metabolism^4,10^. For example, some mutations may be rendered inactive in one metabolic state (a metabolic state is defined by the concentration of each of the metabolites the cell accumulates), while being active in a different one. Switching of metabolic niches may expose certain phenotypes that are hidden by metabolism-dependent genetic buffering. 2) tissue-specific metabolic differences may predispose cell-types to aggregate or misfold particular mutant protein, even while the protein is ubiquitously expressed. Age-dependent change in metabolism may also render certain tissues more prone to age-dependent aggregation^11,12^. This may have implications in explaining the late-onset tissue specificity of aggregation associated disorders^13^ which has been hard to explain with our current understanding of proteostasis components.

Given the immediate importance of the second implication, the link between cellular milieu and proteostasis should be investigated with human diseases in mind^14^. However, given that metabolism is intricately linked to regulation of molecular chaperones through nutrient-signaling (TOR, gcn2, AMPK and so on) it is difficult to pinpoint any proteostasis differences found in eukaryotic models to cellular milieu^15–18^. It will be hard to exclude the role of canonical molecular chaperones, autophagy or translation related mechanisms that are extensively studied^19–21^. Although these systems are extremely important in explaining cellular quality control, the complexity of these machineries renders unmasking the role of cellular milieu in proteostasis a daunting task. In addition, the growth conditions for cell lines (specifically non-cancerous and untransformed cells needed in the study) are hardly defined. This place a limitation in understanding or altering metabolism in defined directions.

In order to test this fundamental link between metabolites and proteostasis, we needed a bare minimum framework, an *in vivo* test tube, to test this possibility. We chose *E. coli* as it is well characterized in terms of its metabolic network, its chaperone content and the network of chaperones that work in the cytosol^22,23^. It also has few defined mechanisms to upregulate chaperones. We use this model and a newly devised assay to address the possibility of metabolism affecting intracellular protein folding.

One of the ways proteostasis can be studied is by monitoring the capacity of cells to buffer non-synonymous mutations^24,25^. While many studies have shown mixed results in terms of buffering by molecular chaperones-the proteins that aid in folding^26–30^, remarkably little is known regarding the role of cellular chemical milieu in proteostasis and mutational buffering. Previously it has been shown that the addition of small molecules at large concentrations in growth media leads to mutational buffering^25,31^. Interestingly, the type of mutations buffered depended on the nature of the small molecule as different small molecules have different mechanisms to aid protein folding^25^. This indicated that molecular evolution can take different routes if different molecules are present in the growth media^32^. However, we do not understand if the physiological concentrations of metabolites present inside the cell affect protein folding and mutational buffering.

Osmolytes are abundant in cells and cells respond to osmotic shock by rewiring metabolism^10,33^ which allows them to accumulate compensatory osmolytes^34^. Osmolytes also influence protein stability *in vitro*^32,35–38^. We hypothesized that change in the osmotic composition of a cell may thus change protein folding, and hence mutational buffering. To test this, we used strains with altered levels of certain osmolytes and monitored their potential to buffer mutations in two model proteins. Indeed, the mutational buffering capacity was altered when the metabolite pools were altered. Interestingly, buffering capacity of the same strain in different metabolic states was different. In all cases, mutational buffering was only evident for mutations that impair folding, corroborating the link between proteostasis and genetic buffering. Remarkably, the metabolites that change along with buffering capacity were able to aid protein folding *in vitro*, suggesting a strong link between metabolite-assisted protein folding and genetic buffering. Finally, we proved the link between metabolic state and mutational buffering by evolving strains of *E. coli* with enhanced osmotic tolerance. These strains showed similar altered buffering capacity as seen for metabolically compromised cells, highlighting that the protein folding environment is different in different metabolic states. We propose that metabolic alterations can have far-reaching consequences on mutational buffering through their influence on proteostasis.

## Results

### Altered metabolite uptake leads to differences in mutational buffering

To elucidate if metabolic rewiring changes capacity to buffer mutations, we used two model proteins-Gentamicin-acetyl transferase (Gm-R, confers gentamicin resistance)^39^ and Green Fluorescence Protein (GFP - yeast enhanced variant)^40^. These proteins met a few essential requirements. 1) Employing these model proteins, we could monitor the activity of multiple mutants simultaneously. 2) These proteins are non-endogenous to *E. coli* and their activity is largely independent of endogenous *E. coli* gene regulatory network except for the proteostasis network that takes care of its biogenesis and degradation. It ensured that altered buffering of different mutants of the proteins in different conditions is indeed an alteration in the general mutational buffering capacity of *E. coli* (**Figure 1**). Endogenous model proteins are attractive in terms of their potential to unearth the native dependence on the *E. coli* environment. However, mutational buffering for these mutants are complicated by buffering pathways that are specific to the protein chosen (**Figure 1**). This is overcome by the use of exogenous proteins. 3) The two chosen proteins are unique protein-folds, presumably with different folding requirements. This enabled us to exclude fold-specific artifacts. 4) GFP was amenable to *in vitro* protein folding studies, allowing us to reconstitute the buffering activities *in vitro* thereby helping us to delineate the molecular mechanism of buffering.

**Figure 1:**
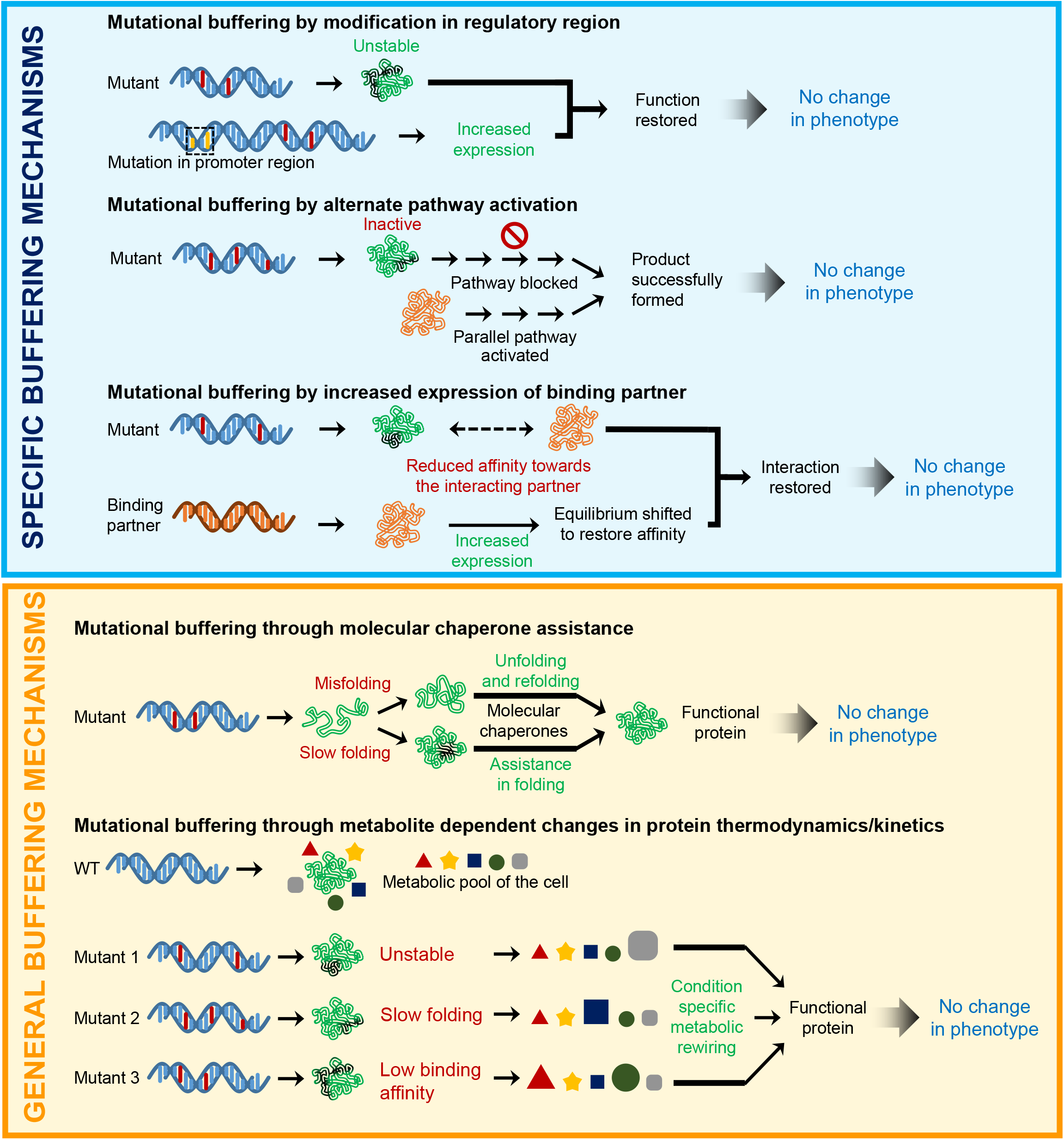
Schematic showing specific and general buffering mechanisms. Mutational buffering in a cell can be achived by modification in the regulatory region of a substrate leading to its increased expression, activation of alternate pathways or by increased expression of the binding partner for a specific substrate (top panel). However generalized buffering of mutant phenotypes is executed by protein folding assistance through molecular chaperones or through relative composition and concentration of metabolite and resultant changes in protein thermodynamics and kinetics (bottom panel).

To generate a comprehensive map of mutational buffering, we developed massively-parallel activity assays to quantitate protein activity of a large number of mutants of the test proteins. Since Gm-R confers resistance to Gentamicin (Gm), we modified and developed a high throughput assay to quantitate protein activity using deep-mutational scanning^24^. We used a Glycine doublet mutant library for this protein^25^ (**Figure 2A and S1A**). For the second test protein GFP, where we could isolate clones with different levels of GFP fluorescence, we used a random mutant library of GFP. Quantification of the intracellular folded fraction of GFP was done using flow-cytometry. Wt mCherry was expressed along with GFP as a bicistronic construct to control for differences in promoter activity and plasmid number (**Figure 2B**).

**Figure 2:**
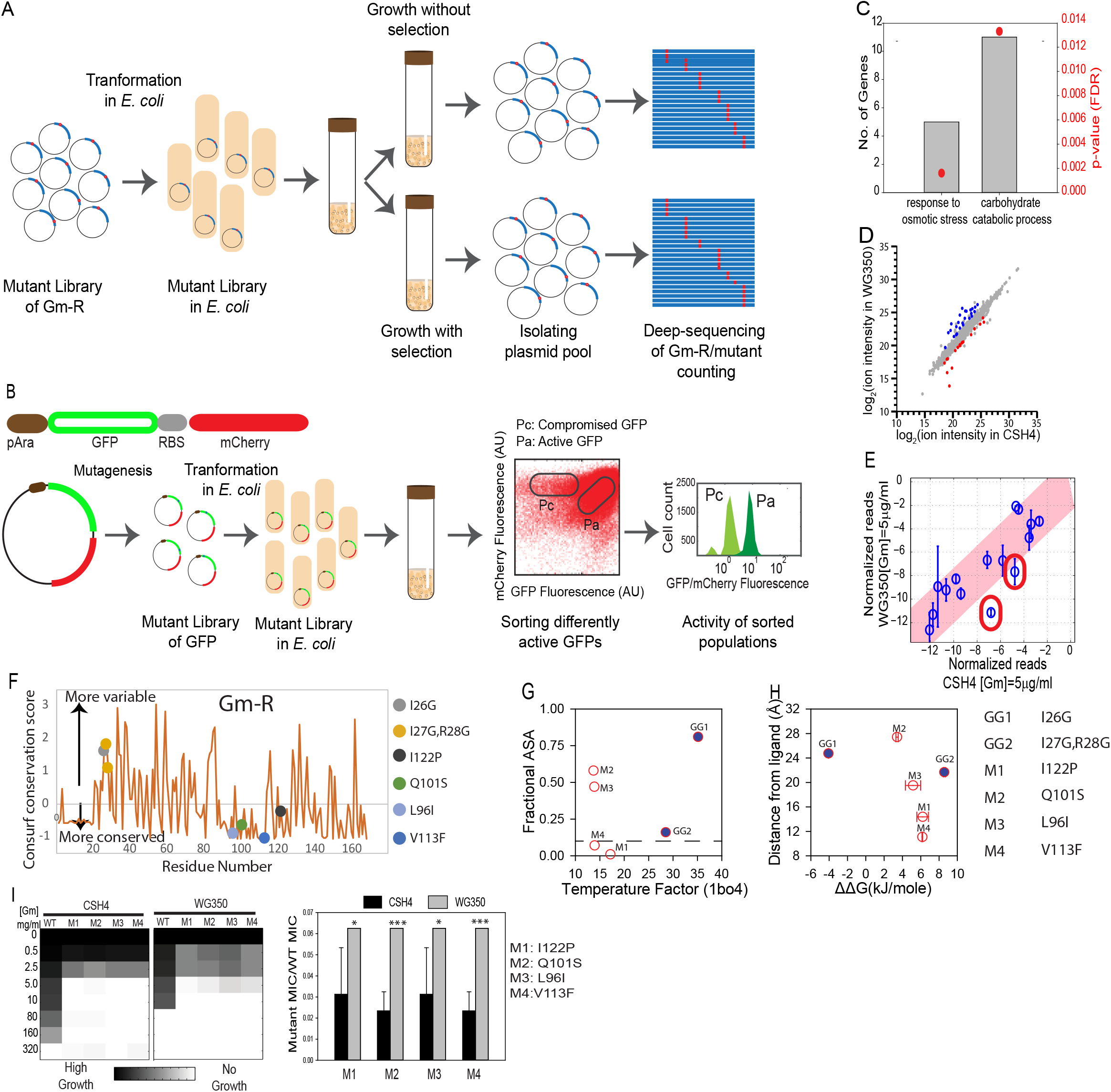
Genetic alteration in cellular metabolism changes mutational buffering. **A.** Schematic of highthroughput activity assay for Glycine-doublet substitution (GG-mutant) library of Gm-R. Activity of Gm-R mutants is inferred from the competitive fitness of the mutants in presence of Gm-selection. The competitive fitness for each of the mutants is quantified by deep-sequencing. **B.** Schematic of highthroughput activity assay for GFP mutant library. The bi-cistronic construct of GFP and mCherry is driven by an arabinose inducible promoter. RBS indicates the position of the additional Ribosome Binding Site for translation of mCherry to serve as internal control. Mutations are created on GFP using random mutagenesis with GFP specific primers. Pool of GFP mutants is sorted into population of compromised mutants (Pc) and active mutants (Pa) based on GFP Fluorescence. **C.** Gene Ontology (GO) classes that were upregulated in WG350 transcriptome with respect to CSH4. The fold enrichment is shown on the left-axis and Benjamini-Hochberg FDR corrected p-values for the enrichment score are shown on the right axis. **D**. Comparison of metabolic features measured in WG350 and CSH4 using untargeted metabolomics. Metabolite features that are significantly different between CSH4 and WG350 are represented as colored circles (p-value < 0.05, 5 biological replicates for each sample). **E.** Normalized read counts of Gm-R GG-mutant library at comparative selective pressures in the presence of the antibiotic gentamicin (Gm). Pink shaded area marks the 99% confidence interval. Mutants marked in red show lower read-counts in WG350 than in CSH4. F. Conservation score (calculated by consurf (Ashkenazy, H. *et al*, 2016)) for each residue. Higher consurf scores indicate more variable residues (less conserved among homologs. Each filled circle indicates the residues that are mutated in the clones studied. G. Fractional ASA (calculated using VADAR (Willard, L. *et al*, 2003)) of the residues mutated in the clones is plotted against the temperature factor of the residues obtained from 1BO4 (left panel). H. Distance of the residues mutated from the ligand was calculated from the pdb (1BO4) and plotted against the predicted alteration in protein stability. Positive ΔΔG corresponds to destabilization and negative to stabilization (calculated by foldX (Schymkowitz, J. *et al*, 2005)). **I.** Left is a heat map representing growth of four Gm-R mutants in WG350 and in CSH4 along with Wt Gm-R in increasing concentration of Gm. Right panel shows the MIC of each of the mutants normalized with respect to Wt Gm-R in the respective strains. Error bars are representative of standard deviation from 4 biological replicates. Significance is calculated using Students’ t-test with respect to CSH4 (* p-value < 0.05, *** p-value < 0.001). **Also see Figure S1**

To obtain *E. coli* strains with altered metabolism, we chose the strain CSH4^41^ and a mutant strain on this background deleted of Proline and Glycine-betaine uptake transporters (WG350) (see Key resource table)^42^. We chose this mutant as it is osmosensitive as compared to CSH4 (**Figure S1B**) suggesting an altered concentration of intracellular osmolytes; this served as an experimental model for alteration in metabolism and osmolyte composition. To validate that deletion of uptake-transporters altered metabolism in the mutant strain we obtained the mRNAs differentially expressed between WG350 and CSH4 using RNA-seq; differentially expressed mRNAs were significantly enriched for genes encoding metabolic enzymes (**Figure 2C**). To substantiate if the strains differed in terms of metabolite concentrations, we obtained an untargeted metabolite profile for both the strains (**Figure 2D, S1C**). The strains differed in terms of some of the metabolic features (43 metabolites with p-value <0.05). For example, the metabolic features corresponding to Trehalose and Trehalose-6-Phosphate increased in WG350 compared to CSH4 by ~3 and ~4-fold, respectively (**Table S1**). Proline and Betaine, although the transporters were deleted in WG350, were only marginally lower (insignificant) in WG350 compared to CSH4 (**Table S1**), indicating the cellular biosynthetic processes are switched on in the absence of transporters. For Glycine-betaine we found a coherent increase in the transcripts encoding genes for its endogenous synthesis and an increase in metabolites involved in its synthesis (**Figure S1D**). This demonstrates that cellular metabolism is rewired significantly upon the deletion of transporters for Proline and Glycine-betaine. Taken together metabolism in CSH4 and WG350 strains were significantly different; this provided us the platform to ask if mutational buffering differed between these strains that differed in metabolism.

To check for mutational buffering, we transformed both the strains with Gm-R Glycine-duplet substitution (GmR-GG) library and grew them at similar selection pressure (see star methods for details) in the presence of Gentamicin (Gm) (**Figure S1E**). To check for mutation-specific effects it was important to normalize Minimal Inhibitory Concentrations (MICs) of the mutants to that of Wt Gm-R, as Wt Gm-R transformed WG350 was more sensitive to Gm than CSH4 with Wt Gm-R (**Figure S1E**). Mutant pools were sequenced and analyzed to the obtain abundance of the different mutants in the presence or absence of selection pressure (Gentamicin in growth medium). Enrichment scores in the presence of Gm selection were calculated to quantitate their activity as previously published^36^. Enrichment scores (a measure of activity) of a chosen set of partially active and inactive mutants as identified with this assay correlate well with the semi-quantitative measurements of activity as obtained by minimum-inhibitory concentration (MIC) of Gm in presence of each of these mutants (**Figure S1F**). While most mutants are similarly active in the two stains CSH4 and WG350, two of the partially active Gm-R mutants (Gm-R (I26G) and Gm-R (I27G, R28G)) were less active in WG350 than in CSH4, indicating mutation-specific differences buffering (**Figure 2E**). Rate of translation/transcription changes protein folding environment. We confirmed that transcription/translation was not different between these two strains using a GFP/mCherry system that is driven by the same promoter (**Figure S1G**). The Gm-R mutations buffered differently between strains were in the non-conserved region of the protein (**Figure 2F**), exposed to solvent, and flexible (**Figure 2G**). One of the mutants was predicted to decrease the stability of the protein indicating folding problems, however, none of them were located near the active site (**Figure 2H**). This suggested that alteration in metabolism may affect the folding of these mutants in WG350 strains. Remarkably, a different set of small-molecule-dependent Gm-R mutants isolated in a previous study^25^ show higher activity (w.r.t Wt Gm-R) in WG350 than in CSH4 (**Figure 2I**). All these mutant proteins harbored mutations in conserved residues (**Figure 2F**), that were relatively inflexible (**Figure 2G**). While these mutations were away from the ligand-binding site, all of them are predicted to partially destabilize the protein (**Figure 2H**). This indicated that mutations even in the buried, conserved residues that destabilize protein structure can be buffered differently in the two strains.

To compare mutational buffering between WG350 and CSH4 for the second test protein with a completely different fold, we purified a population of GFP mutants (compromised fluorescence, Pc) that show lower GFP fluorescence compared with Wt GFP in the WT *E. coli* strain BW25113 (referred to as BW henceforth). The mutant pool (Pc) had similar fluorescence in CSH4 and WG350 strains (**Figure S1H**), demonstrating that there is no difference between the strains in buffering mutations on GFP and that the difference between the strains in buffering mutations is protein-specific. Taken together, this suggests that the ability to take up metabolites from the medium affects metabolic network and mutational buffering. This change the spectrum of mutations buffered in a protein-specific manner.

### Different metabolic states of the same cell show differences in buffering capacity

Next, we asked if altering the metabolite pool in the same strain background changes mutational buffering. Since osmotic shock alters the metabolite pool facilitating the accumulation of osmotically active metabolites^28^, we grew the strains WG350 and CSH4 in 350mM NaCl to change the metabolic status of the cell. We obtained transcriptomic (**Figure 3A**) and metabolomic shifts (**Figure 3B, S2A**) associated with osmotic shock in each of the strains. Osmotic shock altered metabolism in both the strains (**Figure 3C, S2B**) but was routed differently in the two strains. For example, levels of Fructose 1,6-bisphosphate increased in CSH4 upon osmotic shock but not in WG350, contrastingly, Succinate increased in WG350 but not in CSH4 upon osmotic stress (**Table S1**).

**Figure 3:**
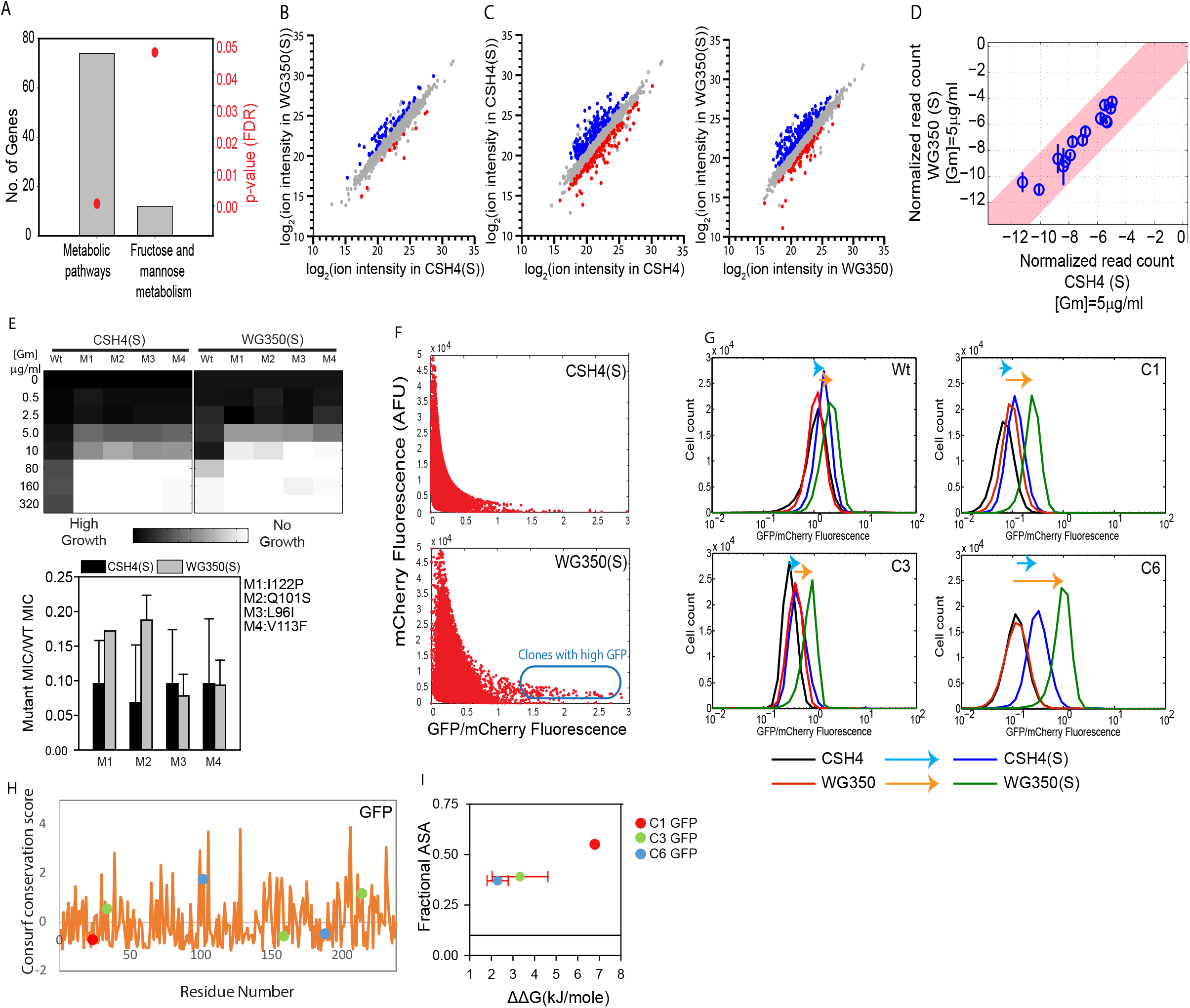
Osmotic stress-induced changes in metabolism changes the spectrum of mutants buffered. **A.** Gene Ontology (GO) classes that were upregulated in WG350 transcriptome with respect to CSH4 during osmotic shock. The fold enrichment is shown on the left-axis and Benjamini-Hochberg FDR corrected p-values for the enrichment score is shown on the right axis. **B.** Comparison of metabolic features measured in WG350 and CSH4 during growth in media containing 350 mM NaCl using untargeted metabolomics. The significantly different metabolites (p-value < 0.05, 5 biological replicates for each sample) are represented in colored circles. **C.** Comparison of metabolic features within strain in presence and absence of 350 mM NaCl in growth media. The significantly different metabolites (p-value < 0.05, 5 biological replicates for each sample) are represented in colored circles. **D.** Normalized read counts of Gm-R GG-mutant library at comparative selective pressures in the presence of the antibiotic Gentamicin (Gm) and 350 mM NaCl in the growth media. Pink shaded area marks the 99% confidence interval. **E.** Top panel shows heat map representing activity (in terms of growth) of Gm-R mutants described earlier in CSH4 and WG350 grown in presence of 350 mM NaCl and different concentration of Gm. Bottom panel shows the MIC of each of the mutants normalized with respect to Wt Gm-R in the respective strains. Error bars are representative of standard deviation from 4 biological replicates. **F.** Single cell fluorescence for the partially active pool of GFP mutants (Pc). The scatter plot for mCherry Vs GFP/mCherry fluorescence is shown for Pc in WG350 and CSH4 when grown in the presence of 350 mM NaCl added in excess to the LB growth medium. Clones with high GFP fluorescence during osmotic stress in WG350 are indicated by the blue box. **G.** Histogram of GFP/mCherry fluorescence of the Isolated clones (C1,C3,C6, and Wt GFP) in WG350 and CSH4 in the presence and absence of 350 mM NaCl. The shift in the ratio due to osmotic shock is shown with colored arrows. The cyan arrow indicates increase/decrease in GFP fluorescence upon addition of 350 mM NaCl to CSH4 strain, the orange arrow indicates the same for WG350. **H.** Conservation score (calculated by consurf (Ashkenazy, H. *et al*, 2016)) for each residue. Higher consurf scores indicate more variable residues (less conserved among homologs. Each filled circle indicates the residues that are mutated in the clones studied. I. Fractional ASA (calculated using VADAR (Willard, L. *et al*, 2003)) of the residues mutated in the clones is plotted against the temperature factor of the residues obtained from 1GFL (left panel). Positive ΔΔG corresponds to destabilization and negative to stabilization (calculated by foldX (Schymkowitz, J. *et al*, 2005)). **Also see Figure S2**

Interestingly, the osmotic adaptation of CSH4 and WG350 strains led to a marked similarity in terms of their potential to buffer di-glycine mutations in Gm-R (**Figure 3D**). Gm-R (I26G) and Gm-R (I27G, R28G), mutants less active in WG350 than in CSH4, show similar activity in these strains in the presence of NaCl in the growth medium. The mutants of Gm-R that showed enhanced activity in WG350 compared to CSH4, shows a noticeably smaller difference or similar activity in both the strains during osmotic shock (**Figure 3E**). This rule out the canonical effect of osmotic stress-induced aggregation in affecting buffering under the conditions used here. Notably, the buffering capacity of WG350 and CSH4 towards Gm-R was similar although the osmotic composition of the strains in the presence of the salt is different. This indicates that compensatory mechanisms may work through different metabolic pathways to reinstate a similar spectrum of mutational buffering.

Since metabolite composition of WG350 and CSH4 were different in the presence of salt, and it is known that different small molecules have protein and mutation-specific effects^32^, we asked if there could be difference in mutational buffering for a different protein. Indeed, mutational buffering of these strains during osmotic shock is different for the GFP mutant library (although for Gm-R it was the same) (**Figure 3F**). We could identify multiple clones of GFP that showed enhanced fluorescence in WG350 compared to CSH4 in the presence of osmotic shock. This clearly shows that the activity of these GFP mutants are enhanced in this altered osmotic condition. In order to validate the observed buffering, we isolated single clones from the pool using FACS and sequenced them. The buffered pool isolated was enriched in mutation G24C (C1 GFP) amongst other mutations E34G, N159K, K214T (C3 GFP) and D102N, I188N (C6 GFP). (**Figure S2C**). Upon retransformation, each of these mutants exhibited similar fluorescence in CSH4 and WG350 in normal growth media while their fluorescence increased in osmotic stress (**Figure 3G, S2D**). Notably, these exhibited not only higher fluorescence in WG350 than in CSH4 in the presence of osmotic shock but different mutants were buffered to different extent in WG350 under osmotic stress. Each of these clones had at least one mutation that mapped onto highly conserved region of the protein (**Figure 3H**), were exposed and predicted to destabilize the protein structure (**Figure 3I**). These clones confirm that differences in metabolism can buffer mutations in conserved regions that tend to destabilize proteins, but in a protein specific manner in different conditions.

### Mutational buffering is affected through altered proteostasis

Having obtained different clones of GFP we asked if mutational buffering has contribution from altered proteostasis. The mutations did not map close to the fluorophore of GFP^43^ (**Figure S3A**) and *in vitro* fluorescence of the purified mutants was like that of Wt GFP (**Figure 4A**) confirming that the mutations did not affect fluorescence of the folded proteins. Refolding studies of the purified proteins showed that the apparent rates for refolding was unchanged for C6 while C1 refolded with a rate ~10 fold slower and C3 refolded with a rate ~5 fold slower than Wt GFP (**Figure 4B, S3B**). However, the native state of C1 and C6 mutants were as stable as Wt GFP towards temperature denaturation (**Figure 4C, S3C**) while C3 was marginally destabilized, suggesting that defect in folding may depend on either the stability of the folding intermediates or that of the native state. This demonstrated that the mutants isolated were indeed folding-compromised mutants, most probably resulting from energetic frustrations in the folding landscape. Thus, differences in their *in vivo* fluorescence in different strains and conditions may reflect the differences in proteostasis capacities between strains.

**Figure 4:**
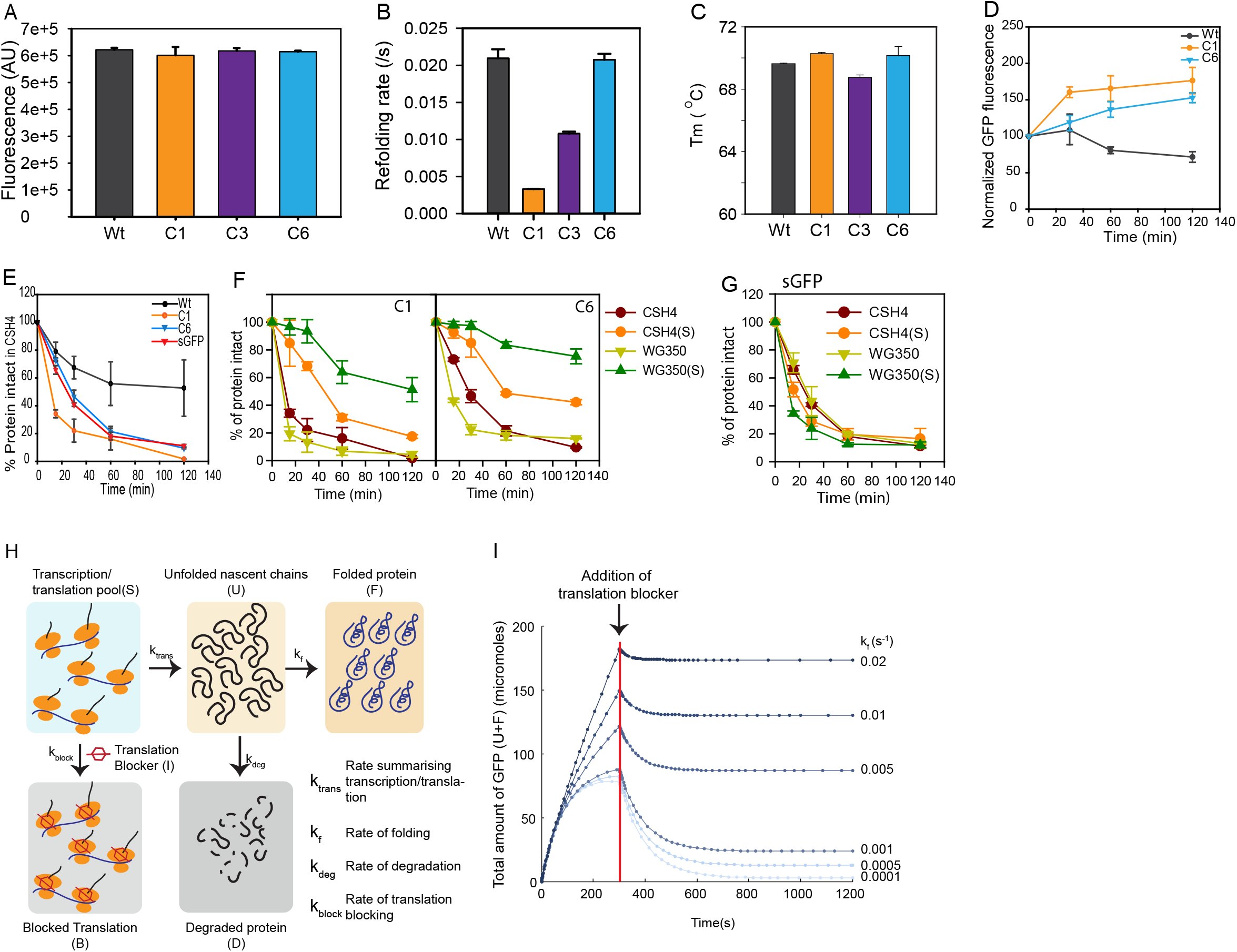
Mutations buffered by altered metabolic states have compromised protein folding. **A.** Fluorescence of Wt GFP and GFP mutants (C1, C3 and C6) at 515 nm are shown at 200 nM concentration (excitation 488 nm /slit-width 2 nm, emission slit-width 5 nm). **B.** Refolding rates of Wt and GFP mutants obtained on unfolding the proteins in 6 M GuHCl, followed by a 100-fold dilution into Buffer-A (50 mM Tris, 150 mM NaCl, 2 mM DTT, pH 7.4) to a final concentration of 200nm for the proteins. Refolding rates were obtained by fitting the refolding traces with single exponential equations. **C.** Average melting temperature (Tm) calculated from thermal melts for Wt, C1, C3 and C6 GFP. Fluorescence of each mutants at different temperatures has been normalized to its respective fluorescence at 25°C. Standard deviations is plotted as error bars from 3 biological replicates. **D.** Plot for Normalized GFP fluorescence of Wt, C1 and C6 GFP at 0 min, 30 min, 60 min and 120 min post translation arrest with Chloramphenicol after 5 hr induction of GFP with 0.5% arabinose in CSH4. E. Plot for the gel-based quantification of intact GFP in Chloramphenicol-chase assay for monitoring degradation of GFP mutants and Wt GFP in CSH4. **F.** Plot for the gel-based quantification of intact GFP in Chloramphenicol-chase assay for monitoring degradation of mutant GFPs C1 and C6 in CSH4, CSH4(S) (CSH4 grown in presence of 350 mM NaCl), WG350 and WG350(S) (WG350 grown in presence of 350 mM NaCl), Standard deviations is plotted as error bars from 3 biological replicates. **G. P** lot for the gel-based quantification of intact GFP in Chloramphenicol-chase assay for monitoring degradation of sGFP in CSH4, CSH4(S), WG350 and WG350(S). **H.** Schematic for simulation setting. Numerical simulation was set up with a fixed concentration of mRNA/ribosome complex (S). These complexes could either form nascent polypeptides (U) with a rate k_trans_ that is a collective rate constant for transcription/translation, or be inhibited with a rate constant k_block_ in the presence of a translation inhibitor (I). The pool of U could either degrade with a rate k_deg_ or fold with a rate constant of k_f_ and reach the native state (F). We finally monitor the total amount of undegraded protein (U+F) after blocking translation after 300 s of starting the simulation (simulation start mimics induction). **I.** Total amount of intact GFP obtained from simulation as a function of k_f_ while keeping kdeg constant at 0.01 s^−1^. Red line indicates the time-point at a 1 mM dose of translation-inhibitor (I) is added. **Also see Figure S3**

To check for *in vivo* folding defects for the isolated GFP mutants, we checked for the degradation rates of a representative set of mutants (slow folding and buffered-C1; fast folding and buffered-C6) vis a vis Wt GFP by stalling translation (**Figure S3D**). First, we checked if the native state of the mutants were less stable than Wt GFP *in vivo* by monitoring fluorescence of the mature protein after blocking translation (**Figure 4D, S3E**). The turnover of the mutants did not differ significantly from that of Wt GFP corroborating with the *in vitro* data that the native state of these proteins is similarly stable *in vivo* at physiological temperatures. Next, to check if folding intermediates of the mutants are less stable than Wt GFP we followed turnover of nascent GFP polypeptides by blocking translation after briefly inducing protein expression (**Figure 4E**). Total GFP level was monitored using immunoblotting. All the mutants degraded faster than Wt GFP underlining that nascent chains of these mutants are folding-compromised under *in vivo* conditions (**Figure 4E**). This clearly demonstrate that the isolated mutants (C1 and C6) have folding intermediates that are less stable than the folding intermediates of Wt GFP resulting in compromised folding pathways *in vivo*.

To check if the difference in proteostasis between WG350 and CSH4 with and without osmotic shock contributed towards differential buffering in the strains, we checked the degradation rates for the nascently synthesized chains of GFP mutants (as described above) in WG350 and CSH4 in the presence and absence of osmotic stress. The degradation rates of the mutant GFPs in CSH4 and WG350 were similar, however, there was a sharp decrease in degradation rates of the C1 and C6 in WG350(S) (WG350 when grown in 350mM NaCl) compared to CSH4(S) (CSH4 when grown in 350mM NaCl) (**Figure 4F (graph), S3F (gel)**). This was not a general decrease in degradation capacity of the cell, as sGFP - a degradation prone mutants of GFP - degraded with similar kinetics in both the stains in the presence and absence of salt-stress (**Figure 4G (graph), S3H (gel)**). Furthermore, the general proteases were upregulated in WG350(S) rather than being downregulated (shown later in **Figure 5A**) arguing against a possible decrease in degradation capacity of WG350(S).

**Figure 5:**
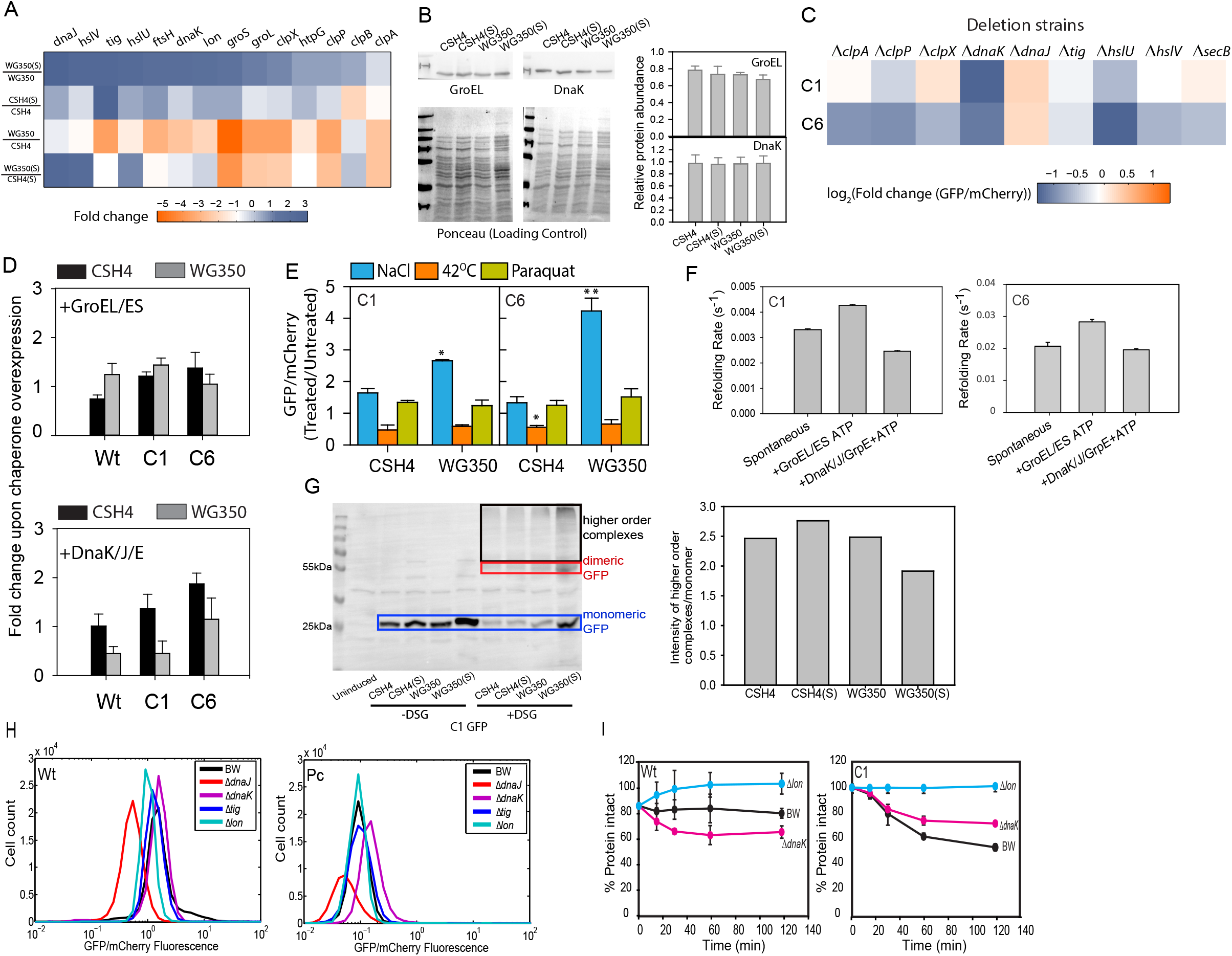
Folding of isolated mutants is independent of molecular chaperones. **A.** Heatmap representing relative quantification of transcripts encoding proteostasis related proteins in WG350 and CSH4 in the presence and absence of 350 mM NaCl. **B.** Left panel shows the immunoblot and right panel the image based quantitation of GroEL and DnaK expression levels in WG350 and CSH4 in the presence and absence of 350 mM NaCl (p-value > 0.05). Ponceau stained blot, used as loading control, is shown at the bottom of left panel. **C.** Wt, C1 and C6 GFP were expressed in wild type *E. coli* (BW25113, BW) and in strains harboring single deletion of different molecular chaperones and proteases. GFP/mCherry for each of the mutants were normalized with respect to Wt GFP fluorescence in the same strains. Fold change in GFP/mCherry for each of the mutant GFPs between BW and the deletion strains was calculated using these normalized values. log_2_ values of this fold change is shown as heatmap. **D.** Wt, C1 and C6 GFP were expressed in CSH4 or WG350 with and without plasmids overexpressing either GroEL/ES (top) or DnaK/J/E (bottom). Median GFP/mCherry fluorescence levels of different strains are shown as a bargraph. Error bars represent standard deviation of 3 biological replicates (p-value > 0.05). **E.** Wt, C1 and C6 GFP were expressed in CSH4 and WG350 in the absence or presence of external stressors (42°C - heat shock, 80 μM Paraquat-oxidative stress, 350 mM NaCl -osmotic shock). GFP/mCherry fluorescence is shown in each of these conditions as bargraph with standard deviations as error bars from 3 biological replicates. Signifance is calculated using Students’ t-test (* p-value < 0.05, ** p-value < 0.01). **F.** To check chaperone dependence of GFP mutants *in vitro*, unfolded proteins were diluted 100-fold in the presence of DnaK/DnaJ/GrpE (400 nM/800 nM/400 nM, respectively) or GroEL/ES (400 nM/800 nM, respectively), and folding was initiated by adding 2 mM ATP. Refolding was followed by monitoring GFP fluorescence as a function of time. Rate of refolding is plotted for C1 and C6 mutants of GFP in presence and absence of chaperoning machinery with error bars from 3 biological replicates. **G.** Immunoblotting for GFP showing the recruitment of nascently formed C1 GFP into multiprotein complexes. After a brief induction of GFP expression, cells were treated with a cell-permeable crosslinker (DSG). Subsequent to crosslinking, cells were lysed and resolved on SDS-PAGE followed by immunoblotting with anti-GFP antibody. Free (or monomeric) GFP is shown with blue box, dimeric GFP is shown in red box and multimeric complexes are shown in black box. The ratio of GFP in multimeric complex to the amount in monomeric state is shown in the accompanying bar graph. **H.** Histogram of GFP/mCherry fluorescence of Wt GFP and pool of GFP mutants having compromised fluorescence (Pc) in BW, molecular chaperone (Δ*dnaJ*, Δ*dnaK*, Δ*tig*) and protease knockout strains (Δ*lon*). **I.** Plot for the gel-based quantification of intact GFP in Chloramphenicol-chase assay for monitoring degradation of Wt and C1 GFP in BW, chaperone (Δ*dnaK*), and protease (Δ*lon*) knockout strain. Standard deviation is plotted as error bars from 3 biological replicates. **Also see Figure S4**

A simple kinetic simulation of protein synthesis followed by modeling competition between folding and quality control assisted degradation (**Figure 4H, Table S2** indicated that an increase in folding rate would be expected to show a decrease in degradation (**Figure 4I**). Thus, the apparent decrease in degradation rate may arise solely due to differences in the folding rate *in vivo* in the different strains and conditions while the degradation rate remains constant (as seen for sGFP). Taken together this indicates that the mutants isolated were indeed folding mutants and mutational buffering differed between the strains due to their differences in proteostasis capacities.

### The mutants are dependent on chemical chaperones and not on molecular chaperones

To investigate the contribution of molecular chaperones, we checked if their levels changed in WG350(S). The set of genes involved in maintaining protein quality control was expected to be induced (or repressed) in WG350(S) than in WG350, CSH4(S) or CSH4. However, mRNA levels of none of the chaperones or proteases other than DnaJ and HslU/V followed the expected over-expression pattern (**Figure 5A**). Protein levels of the canonical chaperones, GroEL and DnaK also did not differ significantly (**Figure 5B**).

Since we isolated specific clones, it provided us with the handle to ask if these mutants were dependent on the canonical abundant chaperones of the cytosol. Single deletions of the abundant canonical molecular chaperones (tig, dnaK, dnaJ, secB)^23^, or the proteases (hslU/V, clpX/P/A)^44,45^, did not decrease fluorescence of these clones significantly (**Figure 5C**) suggesting that the intracellular folding of these mutants were independent of these proteostasis machinery. It would also mean that overexpression of DnaJ or HslU/V seen in WG350(S) at mRNA levels may not contribute to the mutational buffering of the mutants. Coherently, overexpressing GroEL/GroES or the DnaK/DnaJ/GrpE system in CSH4 or WG350 did not increase the fluorescence of these mutants (**Figure 5D, S4A, S4B**), suggesting that the buffering effect in WG350(S) is independent of the concentration of these molecular chaperones. To mimic a global increase in chaperone levels we used other environmental stressors like heat shock and oxidative stress that are known to increase stress-response driven chaperone levels^23,46,47^. The fluorescence of the isolated clones that show enhanced folding in the presence of NaCl induced osmotic stress did not fluoresce better with heat or oxidative stress (**Figure 5E**).

*In vitro*, GroEL/ES machinery could only marginally accelerate the rate of refolding of GFP mutants C1 and C6, whereas DnaK/DnaJ/GrpE did not accelerate the refolding rate of the mutants (**Figure 5F, S4C**). This confirms that these mutants are not the substrates of the abundant chaperone machinery *in vivo* and *in vitro*. To investigate if other molecular chaperones could assist the folding of these mutant GFPs in WG350(S) more efficiently, we studied the physical interaction of these polypeptides chains with molecular chaperones in WG350(S). To check this, we performed *in vivo* crosslinking and measured the amount of higher-order complexes (possibly with components of proteostasis machinery) formed by the mutants GFPs in CSH4 and WG350 with and without osmotic shock (**Figure 5G, S4D**). The levels were not higher in WG350(S), indicating that the chaperone association is not altered in WG350(S). Thus, strain-specific proteostasis differences that alter mutational buffering have contributions from components other than molecular chaperones.

To demonstrate that the mutants buffered in different metabolic states were indeed special in terms of their independence of chaperones and not an artifact of chaperone independence of the model proteins chosen, we checked the chaperone dependence of the partially active (Pc) population of GFP library in molecular chaperone deletion strains (**Figure 5H, S4E**). Indeed, the mutant pool (Pc), as well as Wt GFP, showed decreased fluorescence indicating that Wt GFP itself is recognized by DnaJ. Fluorescence of the mutant pool increased in Δ*dnaK*. This suggested that DnaK can recognize and hold the mutant proteins and prevent their folding. Additionally, we were able to isolate a GroEL/ES dependent mutant of GFP from this pool (Sadat, Tiwari, Chakraborty, Mapa, *paper under review*), proving that the GFP-mutant pool indeed consists of mutants that are efficiently recognized by the endogenous chaperone systems. Thus, the mutant library has mutants that are dependent on the molecular chaperones although these were not differently active in the different metabolic states of *E. coli* we tested. Next, we checked if the isolated mutants C1 and C6 were efficiently recognized by the most abundant chaperone system. Chloramphenicol chase experiment with C1 and C6 in BW and Δ*dnak* and Δ*lon* strain (**Figure 5I, S4F**) clearly shows that the mutant proteins are recognized by the DnaK chaperone system and is effectively degraded only in the presence of DnaK and lon. Taken together this shows that the mutant library of GFP, as well as the mutants identified to be differently buffered in different metabolic states, are recognized and bound efficiently by the endogenous chaperone system. We also compared the activity of the complete Gm-R GG library as well as two mutants (from the same library) in the Δ*dnaK* strain with their activity in BW *E. coli* strain (**Figure S4G**). The pool of mutants, as well as the two isolated mutants, showed a decrease in GmR activity in the absence of DnaK indicating that DnaK assists in the folding of some of the GmR mutants. Thus, the mutant pools (GFP and GmR) tested for mutational buffering contained mutants that were dependent on the endogenous chaperones but the difference in activity of the selected mutants in the different metabolic states was not due to the difference in concentrations of molecular chaperones between the two strains.

Mutant-specificity and protein-specificity of folding assistance *in vivo* are a hallmark of chemical-chaperone mediated folding. To confirm if these mutants were indeed dependent upon chemical chaperones for folding, we obtained their refolding rates in the presence of different metabolites (**Figure 6A**). Many of the metabolites could act as chemical chaperones to accelerate the refolding rates. Molecules like Aspartate, Glycine, chaperoned refolding of two of the mutants (albeit with mutation-specific effect); C6 mutant did not show enhanced refolding rate with any of the small molecules tested. This reiterated the mutant specific effect of different chemical chaperones^25,32^ and suggested that these small molecules could indeed facilitate the folding of these proteins and lead to mutational buffering. Of note, the space sampled in terms of cellular small-molecules was non-exhaustive and no combinations were tried. Importantly, some of the small molecules (like Asp) were also able to increase the thermal stability of the mutants specifically (**Figure S5A**) indicating that these molecules can act in a mutant specific manner to change the energetics of folding. As a control, NaCl or KCl did not have any significant effect on C6 or C3 indicating that salt used for osmotic shock did not affect the thermodynamics of folding of the buffered mutants in general.

**Figure 6:**
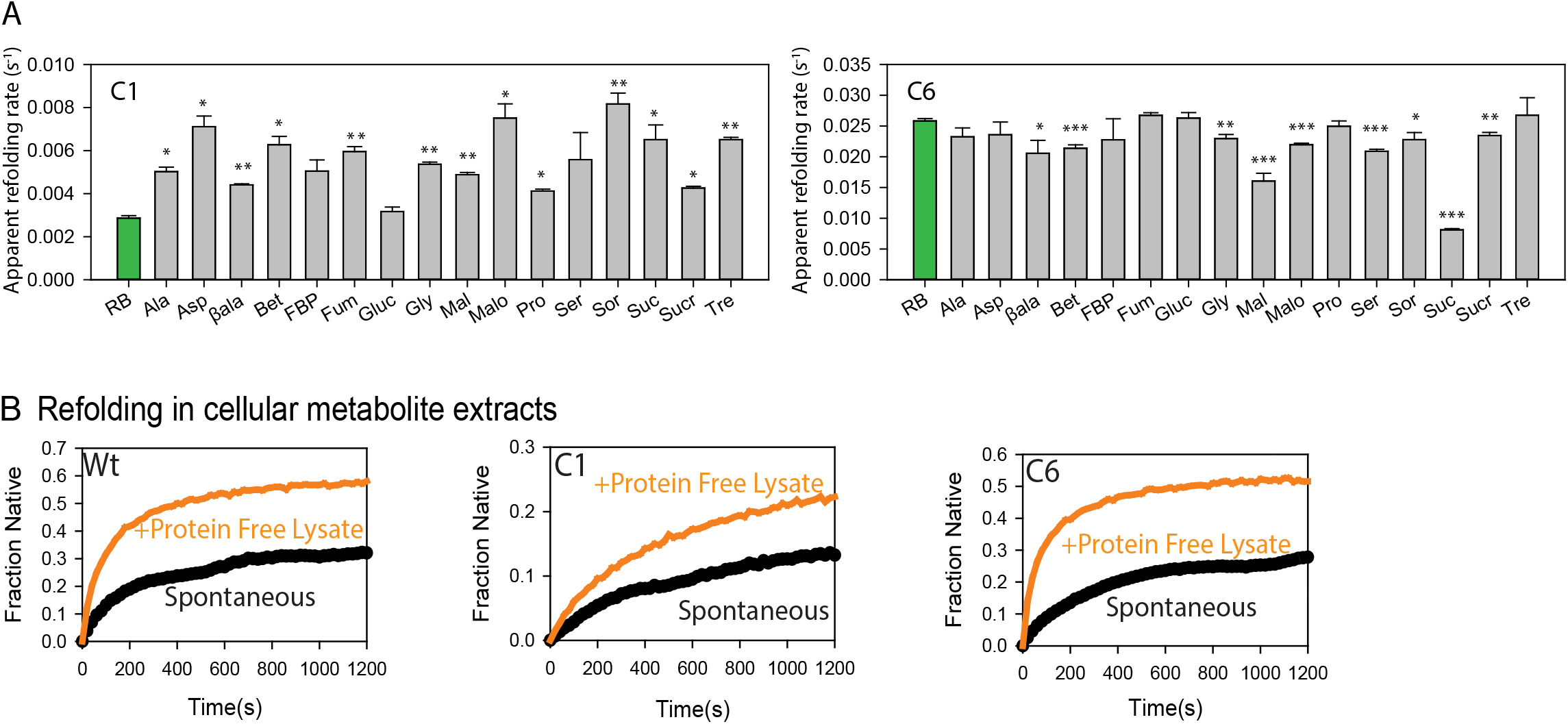
Metabolites can chaperone the mutants *in vivo* and *in vitro*. **A.** Refolding of Wt GFP or mutant GFP was initiated as described earlier in the presence of 100 mM concentration of different metabolites. Refolding rates were obtained by fitting the refolding traces with single exponential equations. Mean of refolding rates, along with standard deviations are shown for three replicates of refolding reactions. RB (shown in green bars) are refolding rates in Buffer-A, the base buffer in which the metabolites are added. p-value is calculated using Students’ t-test with repect to RB (* p-value < 0.05, ** p-value < 0.01, *** p-value < 0.001). Ala: Alanine, Asp: Aspartate, βala: β alanine, Bet: Betaine, FBP: Fructose-1,6-bisphosphate, Fum: Fumarate, Gluc: Glucose, Gly: Glycine, Mal: Malate, Malo: Malonate, Pro: Proline, Ser: Serine, Sor: Sorbitol, Suc: Succinate, Sucr: Sucrose, Tre: Trehalose. **B.** Protein free extracts were isolated from wild type *E. coli* (BW25113, BW) as described in Methods. Refolding was performed by diluting unfolded proteins (Wt GFP or GFP mutants) 100-fold into these extracts.

To check if the small molecule milieu of the cells can aid folding, we reconstituted *in vitro* refolding of the selected GFP mutants in the small-molecule enriched cellular extract of WG350. Refolding of GuHCl unfolded GFP mutants was initiated by diluting out the denaturant in the presence of extract obtained from WG350 (**Figure 6B**). Interestingly, these mutants were refolded to only a negligible extent in the absence of extract while folding substantially in its presence, underlining the importance of the small molecule milieu of the cell in chaperoning protein folding. We confirmed that these lysates were free from proteins and hence molecular chaperones that are known to assist refolding. It is important to stress that the current technologies of extraction limit the concentration at which these small molecules can be extracted; they are more than a thousand-fold diluted from their physiological concentrations. It is therefore difficult to recapitulate the full potential of this mixture and reproduce the *in vivo* differences between strains. However, it demonstrates that the metabolite pool, even on dilution, acts as chaperone and hence can participate in cellular proteostasis. Hence, metabolic differences have the capacity to show altered mutational buffering in different metabolic states. This is primarily evident for mutations that destabilize proteins and make them sensitive to osmolyte concentrations.

Taken together, metabolic differences manifest differences in mutational buffering which may have a significant contribution from metabolite-dependent proteostasis.

### Altering cellular metabolism using metabolites changes mutational buffering

Having realized that genetic and osmotic alteration in metabolism changes the spectrum of mutational buffering, we asked if other routes of altering metabolism may show similar changes in buffering. Addition of excess metabolites - like amino acids-is known to rewire metabolism^48^. We added excess of the amino acids, or sugars and other metabolites to rich media to alter metabolism and check for altered buffering. Exogenous addition of many of these compounds to growth media (individual concentrations provided in **Table S3**) also led to enhanced folding of some of mutants GFP in CSH4 *in vivo* (**Figure 7A**). Specifically, addition of Alanine increased the fluorescence of both the mutant proteins as seen *in vitro* (**Figure 7B**). Further demonstrating that the mutants isolated from the screen predominantly respond to proteostasis alterations due to differences in metabolites, and mutational buffering can be altered by altering cellular concentration of metabolites. Interestingly, different additives had mutant-specific chaperoning activity *in vivo* as seen *in vitro*. This unravels the complex connection between mutational buffering and metabolic status.

**Figure 7:**
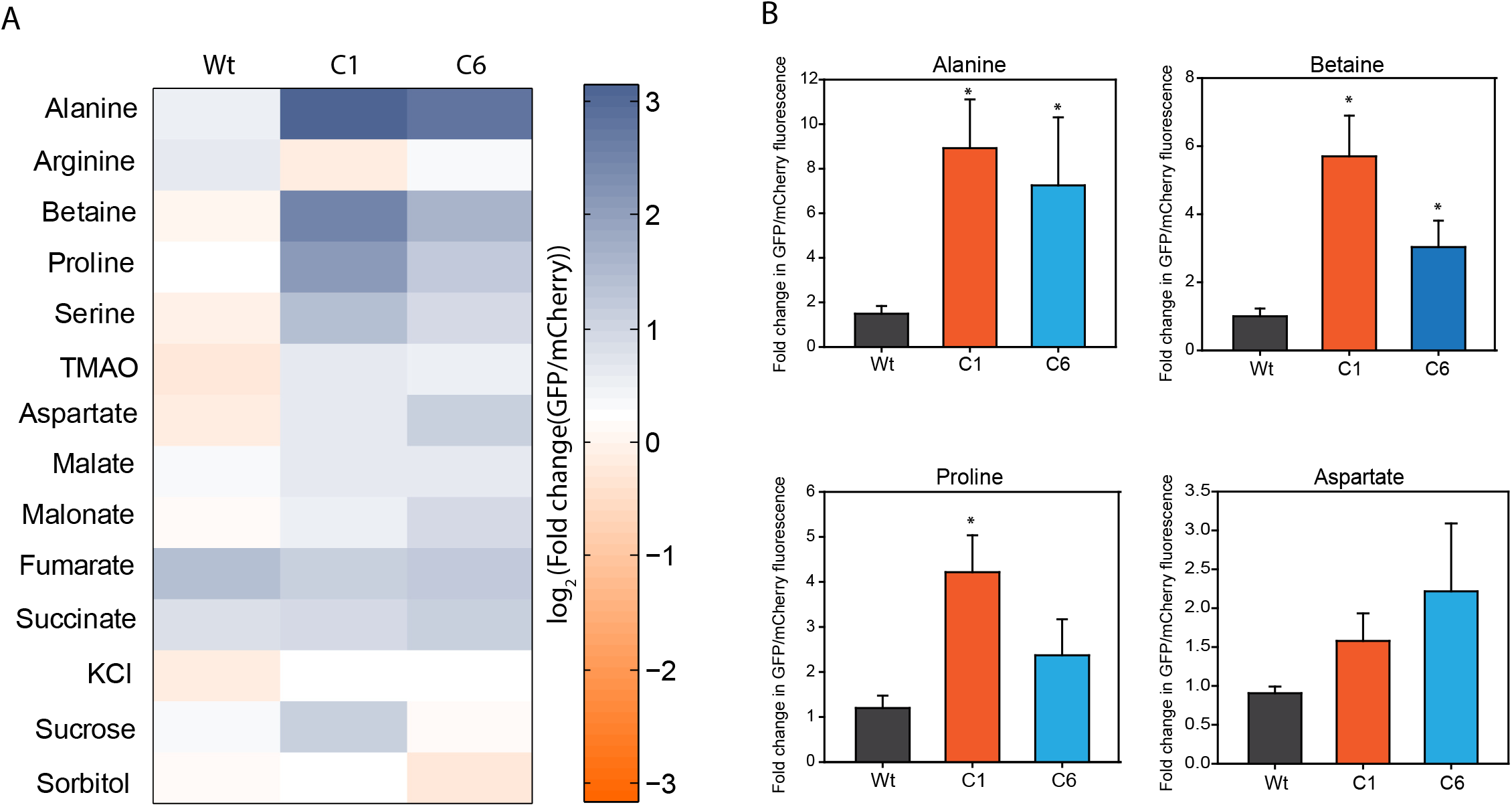
Altering cellular metabolism by exogenous supplementation of metabolites modifies mutational buffering capacity. **A.** Expression of Wt, C1 and C6 GFP mutants was induced in CSH4 cells either growing in LB or in LB containing different metabolites. Log_2_ of fold change in median of GFP/mCherry fluorescence as measured by single-cell fluorescence is shown as heat map **B.** Fold change in median of GFP/mCherry fluorescence in few metabolites (Alanine, Betaine, Proline, Aspartate) in CSH4 is shown as bar graph along with standard deviations from three biological replicates and P-value calulated with respect to Wt GFP by Students’ t-test, * p-value <0.05.

Taken together with our previous observation that these mutants are primarily dependent upon chemical chaperones for folding, we posit a prominent role of metabolism and metabolites in regulating differences in proteostasis capacity.

### Osmotic composition determines the spectrum of mutational buffering

Is the link between metabolism and mutational buffering relevant in the context of natural evolution? To answer this, we checked if the adaptation of an organism in a different niche (that changes metabolism) is associated with the evolution of an altered buffering capacity; we chose to evolve WT *E. coli* (BW25113) to adapt to high salt stress (**Figure 8A**). The strain was continuously passaged in LB in the presence of 500mM of NaCl. We kept the evolution duration short in order to ensure that drift mutations are minimal. Multiple strains were generated by parallel laboratory adaptive evolution, and we obtained strains that were fitter in the presence of osmotic stress by approximately 500 generations (**Figure 8B**). We argued that adaptation for growth in chronic hyperosmotic stress would have fixed mutations that perturb the osmotic composition and hence rewired the metabolic status of the cell. Genome sequencing of two of these strains demonstrates that the each of these strains have accumulated multiple different mutations (**Figure 8C, Table S4**) including mutations in genes involved in osmolyte synthesis (sdaA (gluconeogenesis), fucI (fructose and mannose metabolism), otsA (trehalose metabolism)). This indicated that the strains may have fixed certain traits associated with osmotolerance by acquiring mutations. We confirmed that these strains indeed have a difference in metabolite pool even when grown in the absence of osmotic shock (**Figure 8D**) we could now ask if these metabolic differences changed the mutational buffering capacity. Remarkably, the strains tested showed buffering for the C1 and C6 mutants of GFP that were buffered in WG350 during osmotic stress (**Figure 8E**). The activities of the mutants were two to four-fold higher in the different evolved strains in a mutant-specific manner though the folding of Wt GFP was not altered in these strains (**Figure 8E**). However, *E. coli* molecular chaperones DnaK and GroEL were not upregulated in the evolved strains compared to the unevolved BW strain as checked by immunoblotting (**Figure 8F**) indicating that canonical hubs of proteostasis were not altered in these evolved strains. We confirmed by chase assay that degradation of the buffered mutants was indeed slower in strains that show buffering than in the unevolved strain (**Figure 8G (graph), S6B(gel)**). We confirmed that slower degradation of these mutants in the evolved strain was independent of the activity of the degradation machinery as another degradation prone mutant of GFP (sGFP) degraded as rapidly in the evolved strain as in the unevolved strain (**Figure S6C**); impaired degradation hence is a fallout of faster folding of the buffered GFP mutants in the evolved strains. This indicated that a genetically different strain evolved to have altered osmotic composition is able to buffer similar mutational variation as seen in the WG350 strain during osmotic shock. Response to osmotic shock, or a similar metabolic state once fixed in the genome in the absence of osmotic shock, can buffer similar mutational variations. Like GFP mutants, mutants of Gm-R also exhibited higher activity in most of the evolved strains than the unevolved strain (**Figure 8H, 8I**). Importantly, each of the mutants had different activities in the different osmotolerant strains. Taken together the metabolic state of a cell is directly linked to their ability to buffer mutational variations. Adaptation to a niche with different osmolarity changes the spectrum of mutations buffered. This, in turn, could change the route of molecular evolution of proteins, linking metabolism to the evolution of protein sequences through alterations in proteostasis.

**Figure 8:**
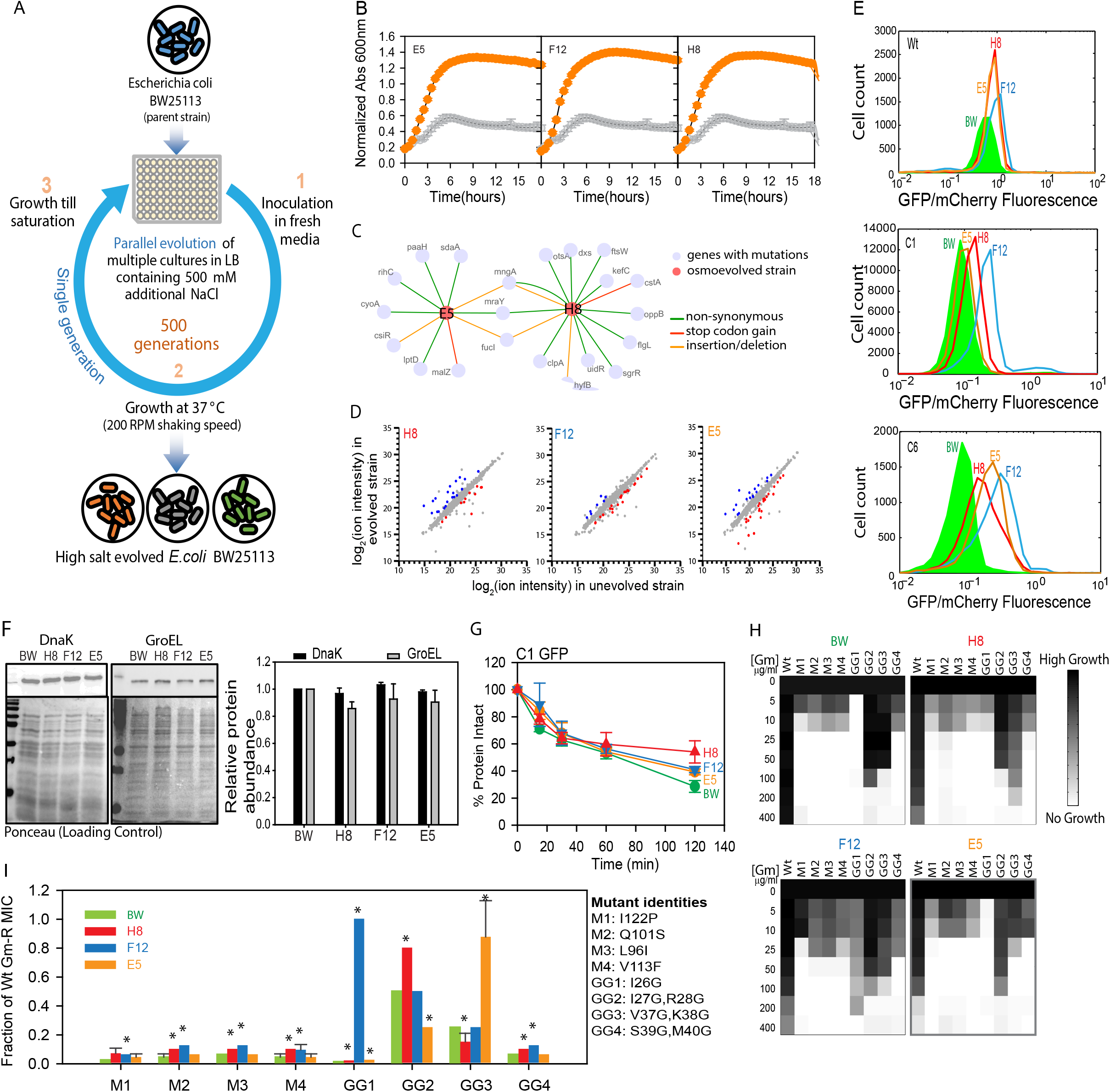
Wild-type cells evolve altered buffering capacity with evolution of a different metabolome. **A.** Schematic of strategy for Laboratory Adaptive Evolution of osmotolerant strains of *E. coli* starting from BW (wild type *E. coli* K-12, BW25113). **B.** Growth curve of unevolved BW and evolved osmotolerant strains E5, F12 and H8 in 500 mM of NaCl added in excess to LB medium while growing at 37°C, 200 rpm. **C.** Genetic interaction network map of evolved strains E5 and H8 based on mutations obtained in genome sequencing. **D.** Metabolite features of representative evolved osmotolerant strains (E5, F12 and H8) and its comparison with BW. The colored circled represent metabolites that are significantly altered in the evolved strains. **E.** Histogram for GFP/mCherry fluorescence of Wt, C1 and C6 GFP in the BW and evolved strains (E5, F12 and H8). **F.** Comparative quantification of chaperone proteins DnaK and GroEL in BW and evolved strains (E5, F12 and H8) done by immunoblotting with specific antibodies as described earlier. Quantification of the chaperone levels are shown as bar graphs. Error bars represent standard deviation of 3 replicate measurement (p-value > 0.05). **G.** Chloramphenicol based chase for checking protein degradation rates were performed in BW and evolved strains (E5, F12 and H8) as discussed earlier. The levels of C1 GFP at different time points are shown as a function of time in different strains. **H.** Heatmap for growth based activity of Gm-R mutants and Wt Gm-R in unevolved strain (BW) and evolved strains (E5, F12 and H8) in increasing concentrations of gentamicin. **I.** Bar graph representing percentage MIC of a selected set of Gm-R mutants w.r.t Wt Gm-R in the respective strains. Error bars represent standard deviation from 2 biological replicates. **Also see Figure S5.**

### Implications

In this work we have indicated the capability of *E. coli*, a prokaryotic unicellular organism, in affecting mutational buffering through accumulation of small molecules in the cell. This, to the best of our knowledge, is a maiden report that specifically describes the chaperoning activity of the small molecule component of the cytosol of *E. coli*. While this fact is important, its regulation through osmotic shock or genetic alterations makes it even more interesting in the context of regulation of protein folding. Regulation of protein folding capacity enables condition specific canalization or decanalization of mutants^49–52^. It is known that perturbing growth conditions by exogenous supplementation by osmolytes can alter mutational buffering, specifically for mutations that affect either protein stability or protein functions^24^. Similarly, endogenous alteration in production of osmolytes (that are part of the metabolite pool) may also change mutational buffering. This we believe would be important when organisms shift niches and evolve new functionalities. It is tempting to speculate that conditions that have been reported to lead to altered Hsp90 levels during the evolution of *A. mexicanus* cave dwelling variants, may also have led to alteration in intracellular osmolyte composition^53^. While *E. coli* is a primitive model organism to comment on adaptive strategies and genetic buffering in higher eukaryotes with complex developmental pathways, it surely paves way for more interesting investigations to establish the potential link between osmotic composition, and their alterations with canalization.

Given that many of the small molecules that accumulate in cells like Arginine^32^, glycerol^54–56^, amino acids^32,57,58^ are reported to change protein folding kinetics and thermodynamics in vitro, it may be the complex milieu of metabolites and not osomolytes alone that can assist or alter the folding process in a protein specific manner. Although worked on for decades, we are yet to understand the molecular basis of protein stabilization by different osmolytes and metabolites. Their mechanisms seem to depend on the test protein, as well as on the specific type of problems that mutants encounter while folding^32^. While many of these molecules (like amino acids and sugars) ^59,60^ can stabilize certain proteins by decreasing flexibility of non-native states, others may prevent formation of non-native contacts or intermolecular protein aggregation by partially destabilizing folding intermediates. Their contribution to cellular protein folding would require further investigation given our observations.

Alteration of metabolic states, as shown in this work provides a new avenue for alteration of proteostasis. Use of a simple model organism serves as an *in vivo* test tube to propose and test the hypotheses related to cellular milieu. In our case we tested that composition of cellular milieu alters with metabolism, and has the potential to alter protein folding and proteostasis network. This link metabolism to protein folding in prokaryotes and imply that microbial evolution may be shaped differently when metabolism is altered either due to environmental pressure of due to genetic changes. Further work will be required to understand if similar mechanisms are prevalent in higher organisms.

## Acknowledgement

The work was done in KC Lab with the grant DST/SJF/LSA-01/2015-16 as Swarna Jayanthi Fellowship Grant from DST, and partly from BSC0124 from CSIR. Work in DS lab was funded through BSC0124 project from CSIR. Instrument support was also obtained from Welcome Trust-DBT India Alliance (KC lab) and CSIR (DS, KC lab). KC acknowledges CSIR and CSIR-IGIB for infrastructural support. KV, RD, AC, MR, ZZ acknowledge CSIR; KS and RD acknowledge UGC; AS acknowledge ICMR, for fellowship support.

## Author Contribution

Conceptualization: Kausik Chakraborty

Supervision: Kausik Chakraborty, Dhanasekaran Shanmugam

Reagent Generation: Kanika Verma, Kanika Saxena, Rajashekar Donaka

Experiments: Kanika Verma, Kanika Saxena, Rajashekar Donaka, Manish Rai, Aseem Chaphalkar, Anurag Shukla, Zainab Zaidi

Analysis: Kanika Verma (FACS/Image/Metabolomics/Sequencing), Kanika Saxena (Mutant library design/FACS), Anurag Shukla (Metabolomics), Manish Rai (Amplicon sequencing/transcriptomics), Rohan Dandage (amplicon sequencing), Dhanasekhran Shanmugam (metabolomics).

Manuscript writing: Kausik Chakraborty and Kanika Verma with inputs from all the authors. All authors read and edited the manuscript.

## TABLES

None

## METHODS

### EXPERIMENTAL MODEL AND SUBJECT DETAILS

#### Strains, Plasmids and Proteins

*E. coli* strain DH5α was used for cloning, WT *E. coli* (K-12 BW25113 referred to as BW) strains was used for expression of arabinose inducible pBAD GFP mCherry and BL21 (DE3) was used for protein expression and purification. Protein concentrations were determined spectrophometrically at 562 nm using BCA kit (Pierce ThermoFisher Scientific). Deletion strains were obtained from CGSC as part of Keio collection^41,42,45^.

### METHOD DETAILS

#### Construction of mutant GFP mutant library

Mutant GFP library was made in arabinose inducible pBAD vector using random mutagenesis approach. In the 1^st^ step YeGFP (referred to as GFP) was amplified using GFP specific primers and Mutazyme II polymerase (Agilent Technologies; Cat no. 200550) in order to incorporate 7-11 mutations per kb of plasmid. In the 2^nd^ step, the product of 1^st^ amplification was used as mega-primer to amplify the entire plasmid with Wt GFP (pBAD GFP mCherry) as template using Kapa Biosystems Hifi readymix (NC02955239) for 25 cycles with both annealing and extension at 72°C. The Wt copy of plasmid was digested with DpnI followed by transformation of chemically competent DH5α cells. The colonies were scraped and plasmid was prepared in pool to yield library of GFP mutants. The said library has a total complexity of around 10,000 mutations. The reporter is constructed such that GFP and mCherry are under same arabinose inducible pBAD promoter in an operon to give readout of GFP according to the mutation created on it but the mCherry readout will remain similar thus serving as an internal control for transcription, translation and inducibility.

#### Screening of mutant GFP library for folding mutants responsive to osmotic stress

WT *E.coli* cells (BW) were transformed with mutant GFP library maintaining 10 fold converge for preserving mutant library complexity. Cells were induced with 0.1% arabinose at the time of inoculation and fluorescence was observed on BD LSR II five hours post induction at 37°C after diluting cells in 1X PBS and incubating at 37°C for 1 hour. Fluorescence of the mutant library was studied in a pooled manner against Wt GFP. The entire library was sorted using BD Aria III into populations of compromised mutants (low fluorescent) and active mutants (high fluorescent) according to the GFP fluorescence. Each of these populations was purified and plasmids prepared. CSH4 and WG350 strains were transformed with the purified populations and subjected to osmotic stress with 350mM NaCl. The pool of mutants buffered under osmotic stress in WG350 was sorted and single clones of GFP were picked from here and checked for their fluorescence in presence and absence of osmotic stress. The isolated mutants having higher fluorescence in WG350 under osmotic stress were identified by Sanger’s sequencing, also cloned in pET SUMO under BamHI and HindIII restriction sites and purified using *E. coli* BL21 (DE3) for further characterization.

#### Growth curve

Single colony of *E.coli* cells were inoculated in LB and grown overnight at 37°C, 200 rpm. Secondary inoculations were done in LB (control) and LB containing 350mM, 500mM NaCl additional in honey comb plates with temperature maintained at 37°C, 200 rpm shaking. Absorbance at 600nm was measured every 30 minutes using Bioscreen C (Oy growth curve Ab Ltd).

#### Transcriptomics

3 ml of culture grown in LB overnight at 37°C from a single colony was used to re-inoculate at 0.1% culture in 10 ml of LB and grown till OD600 reaches 0.5. The culture was mixed with equal volume of bacteria RNA protect reagent followed by RNA isolation using Qiagen RNeasy Mini Kit and TURBO DNA-free kit (AM1907). Quality of RNA was checked using RNA 600 Nano bioanalyzer kit and MICROB Express bacterial mRNA enrichment kit (AM1905) was used to remove rRNA. 100 ng RNA was used to prepare library using Ion Total RNA-seq kit V2 (4475936) and Ion Xpress RNA-seq barcode 1-16 kit. Final quality check and quantification was done using DNA HS bioanalyzer (5067-4626) and qubit HS DNA kit (Q32854). Equal amount of each sample was pooled followed by emulsion PCR (Ion PI Hi-Q OT2 200) and sequencing (Ion PI Hi-Q sequencing 200 kit) FastQC-tool kit was used for Data QC followed by Trimmomatic^61^ to remove low quality reads. TPM was calculated using Kallisto^62,63^ followed by identification of differentially expressed genes using EB sequencing analysis pipeline (https://bioconductor.org/packages/release/bioc/html/EBSeq.html)^64^.

#### Metabolomics

3ml of LB was inoculated with 0.1% inoculum from overnight grown culture and grown till OD600~0.8 at 37°C, 200 rpm. Cells equivalent to 0.1 OD were harvested at 14000 rpm, 1 min, 4°C. Supernatant was discarded and 200ul 80% chilled methanol (80% MetOH, PIPES 1ng/μl, U13C-U15N-glutamine in MS grade water) was added to the pellet and incubated in ice for 5 minutes for quenching. This is followed by sonication in water bath for 15 min at 4°C with intermittent vortexing. Metabolites were collected in the supernatant by centrifugation at 14000 rpm for 1 min at 4°C. Previous step was repeated twice by adding 100μl 80% chilled methanol each time to increase metabolite yield and stored in −80 refrigerator till further analysis. The untargeted mass profile (or metabolic profile) was acquired in Thermo Q-exactive Orbitrap coupled with Thermo Accucore RP C18 150*2.1, 2.6μM column with flow rate of 200μl/min. Ion masses from 80-1000m/z was collected in negative ion mode. The raw output files obtained from mass spec were converted into .mzXML files using proteowizard tool (http://proteowizard.sourceforge.net/) followed by identification and analysis using metabolomics data processing software platform XCMS and MAVEN(http://genomics-pubs.princeton.edu/mzroll/index.php?show=index)^65,66^.

#### Minimum Inhibitory Concentration (MIC) study of Gm-R glycine doublet mutants

The library of Gm-R glycine doublet mutants as reported previously (Bandyopadhyay *et al*., 2012) were grown overnight in LB containing ampicillin (100μg/ml) and Arabinose (0.1%) at 37°C, 200 rpm. Secondary inoculations were done in LB containing Ampicillin (100μg/ml), Arabinose (0.1%) and increasing concentrations of gentamicin (0-800 μg/ml) incubated for 16hrs at 37°C and 200rpm. Growth was assessed by measuring absorbance at 600nm in flat bottom 96 well microtiter plate using TECAN infinite 200 pro. Absorbance value for each sample under selection pressure by gentamicin was normalized against absorbance of respective unselected sample. To obtain a semi-quantitative indication of the activity of different mutants, Wt Gm-R transformed cells were grown as control, and the MIC for the mutants was normalized with respect to the MIC of Wt Gm-R in the same strain.

#### Amplicon sequencing

Plasmids were isolated from unselected (no gentamicin) and selected cells under gentamicin selection pressure. Gm-R gene was amplified for 25 cycles using Kapa Hifi Hotstart polymerase. The 500bp amplicons were gel purified and quantified using qubit. 150ng of purified products was used for library preparation Ion Plus Fragment Library Kit (part no.4471252) and Ion Xpress Barcode Adapters (4471250, 4474009). The final library was quantified and equal amount of DNA library of 8 samples were pooled and emulsion PCR and sequenced using Ion PGM Hi-Q OT2 Kit (part no A29900) and Ion PGM Hi-Q sequencing kit (part no. A30044) on Ion Torrent platform. Analysis was done using FastqQC followed by trimmomatic (cutoff Q15) and quality check by fastQC. Good reads were aligned to Gm-R gene followed by variant calling and fitness score calculation www.github.com/kc-lab/dms2dfe^24^.

#### Quantitation of activity of Gm-R Glycine-doublet substitution mutants

To quantitate activity of the mutants, we obtained the read count for each of the mutant and normalized it with respect to the coverage at the respective positions. The relative read-counts of the mutants did not differ between the different strains in the absence of any selection pressure for Gm-R indicating that the library was homogeneously covered in all the transformations. We thereby did not obtain relative enrichment but compared the coverage-normalized read-counts between the different strains and conditions to obtain differences in activity. All the amplicon-based sequencing experiments were done in duplicates.

To obtain relative activity of the different mutants in one strain (for comparison of activity of mutants as measured by amplicon sequencing and MIC assay as shown in **Figure S1F**) we obtained the enrichment score for each mutant by dividing the normalized read count for each mutant in presence and absence of Gm selection. z-scores were obtained for each of the mutant assuming a normal distribution. A positive z-score indicates higher than average activity while a negative z-score indicated a lower than average activity. Mutants were picked that had a high z-score (1.4, more active than average) or low z-score (−1.6 less active than average) or one close to zero (z-score = −0.4, activity close to average) for checking the MICs. Note: z-score was not used to obtain the significance of difference but the mutants that exhibited either high or low activity.

#### Chase for degradation of protein

From overnight grown culture in LB containing Ampicillin (100μg/ml) secondary cultures were inoculated with 0.1% inoculum in 100 ml LB containing Ampicillin (100μg/ml) and grown till OD_600_ 0.5 or for 5hrs (with inducer for chase of folded protein as shown in **Figure S3E**). Cells were harvested and resuspended in 10ml of the spent media (or media containing 350mM NaCl added in excess to LB. GFP was induced using 0.5% arabinose for 5 minutes at 37°C and 200 rpm. Chloramphenicol (50μg/ml) was used to arrest translation. 1 ml of culture was taken out for uninduced, 0 min, 15 min, 30 min, 60 min, 120 min post translation arrest, harvested, snap freeze and protein was collected by lysing cells resuspended in 1X PBS using 2X lysis buffer (Tris pH 6.8 (80mM), SDS 1%, glycerol 10%). 30μg of protein estimated using Pierce™ BCA Protein Assay Kit and loaded on SDS-PAGE followed by Western blotting probing for GFP using Rabbit anti-GFP (Ab290) and HRP conjugated goat anti-rabbit IgG (SantaCruz Biotechnology).

#### *In-vivo* crosslinking

100ml of LB medium (with and without 350mM of NaCl) containing Ampicillin (100μg/ml) was inoculated with overnight grown cells at 0.1% inoculum for each strain. Cells were grown till they reach O.D_600_ 0.5 and harvested at 4000 rpm for 10 mins at RT. Pellet was finally resuspended in 2ml of spent media and induced for 5 minutes at 37°C with 0.5% arabinose for GFP expression. Cells were harvested and resuspended in 2 ml of 1X PBS containing protease inhibitor cocktail (Roche) and 500μl of resuspended cells was taken as a control for uncrosslinked sample (referred to as -DSG in the text). To the remaining cells, 300μM of Di (N-succinimidyl) glutarate (DSG) was added and incubated at 37°C for 10 minutes for crosslinking. Reaction was quenched with 100mM of Tris, pH 8 for 5-10 minutes at room temperature. Cells were harvested at 13000 rpm, 2 minutes at room temperature and lysed by Freeze-thaw method in the presence of protease inhibitors. 30μg of protein was loaded on SDS-PAGE followed by Western Blotting probing for GFP.

#### Western Blotting Experiment

*E. coli* cells were inoculated in LB and grown at 37°C for 5 hours. Cells were harvested and resuspended in 1X PBS (Himedia; ML023). Equal volume of 2X lysis buffer was added and boiled at 95;C for 15 minutes followed by spin at highest rpm for 15 minutes. The supernatant was collected and estimated for the amount of protein. 30μg of protein was used for SDS-PAGE and Western Blotting post transferring onto nitrocellulose membrane (Millipore; HATF00010). The blots were decorated with antibodies against GroEL (Enzo; 9A1/2) and DnaK (Enzo; 9E2/2) isolated from mouse. Blots were developed using HRP conjugated Goat anti-mouse IgG (Genscript) and luminata crescendo (Millipore). Densitometric analysis was done using ImageJ. (Rasband, W.S., ImageJ, U. S. National Institutes of Health, Bethesda, Maryland, USA, http://imagej.nih.gov/ij/, 1997-2011).

#### GFP refolding assay

60μM native GFP (or mutant) was mixed with 8M GuHCl prepared in GFP refolding buffer (Buffer-A) (Tris-Cl (25 mM), KCl (150mM), MgCl_2_ (10mM), DTT (2 mM), pH 7.4) in 1:3 ratio. After incubation at 25°C for 1 hour, refolding of GFP was initiated in Buffer-A by diluting the mixture 100-fold (final GFP concentration 150 nM). Real time fluorescence of GFP was monitored in Fluorolog 3 spectrophotometer (Horiba Jobin Yvon, with operating software FluorEssence v3.0) at 25°C, with excitation wavelength 488 nm (2 nm slit width) and emission wavelength 515 nm (5nm slit width), enabling ‘anti-photobleaching’ mode. The data acquired from refolding assays was analyzed in OriginPro 8 (OriginLabcorporation).

#### Thermal melt

Buffer-A with or without 100 mM small molecules was added with purified GFP (Wt or mutant) to a final concentration of 150nM. 20μl of this whole mixture was aliquoted in a 386 well RT-PCR plate. The thermal melt was carried out in CFX384™ Real Time System attached to a C1000 Touch Thermal Cycler (Biorad). The program was set to go from 25°C to 90°C with a step of 1°C and incubation time of 2 min/°C. The data was collected as GFP fluorescence at each step, and fit to a linear equation to obtain midpoint of each curve as melting temperature (Tm).

#### Isolation of protein free cell extract

*E. coli* K12 (BW25113) strain was grown in 5 ml LB broth at 37°C, 200 RPM orbital shaking for overnight and this primary culture was used to inoculate a 500 ml secondary culture in LB broth. After growing the secondary culture in identical conditions for 3 hours (OD 0.7-0.8), the culture was centrifuged at 4000 RPM for 30 minutes at 37°C. 5 ml of boiling hot ultrapure water was added directly to the pellet, resuspending it vigorously, and the cell suspension was immediately collected in a glass test tube. The glass tube was kept in a boiling water bath for 30 minutes, ensuring maximum lysis of cells and precipitation of proteins. The resulting suspension was cooled down to room temperature and was centrifuged at 13000 RPM for 20 minutes. Absence of protein in the supernatant, the protein free cell extract, was confirmed by Bradford protein estimation assay.

#### Simulation

Numerical simulations were performed using the ODEs defined in **Figure 4** to model a basic framework to check the apparent rate of degradation as a function of folding rate in vivo. The simulation was set up with 1000μM S (pool of DNA/RNA that is competent to make proteins). S can form U (nascent polypeptides) with an overall unimolecular rate constant of ktrans (a simplistic combination of rates of transcription and translation). The pool of U can either convert to F (folded GFP) or be degraded with the unimolecular rate constants of kf and kdeg, respectively. The pool of S can be blocked by I (inhibitor of translation) with a rate constant of kI. This was included to mimic translation arrest by chloramphenicol. The simulation was run in the absence of I for 300 seconds. Following this, 1mM of I was dosed into the simulation to rapidly quench S. Following this we monitored the total concentration of uncleaved GFP (U+F) over time to mimic the results obtained from the chase experiments performed with anti-GFP antibodies.

#### Conservation score and fractional ASA measurement through FoldX

PDB structures were used to calculate the conservation score of each of the residues using multiple sequence alignment through CONSURF^67^. Fractional ASA, and temperature factor of a residue was obtained using VADAR^68^. FoldX package was used to obtain the predicted stability value for each of the mutants^69^. The calculations were repeated at least 3 independent times to obtain an average and standard deviation value for the predicated stability values.

### QUANTIFICATION AND STATISTICAL ANALYSIS

Student’s t test and R package for non-linear regression was used for statistical analysis. Flow-cytometry data was analyzed using octave

### DATA AND SOFTWARE AVAILABILITY

All data are provided in the manuscript. We did not develop any new software.

## KEY RESOURCES TABLE

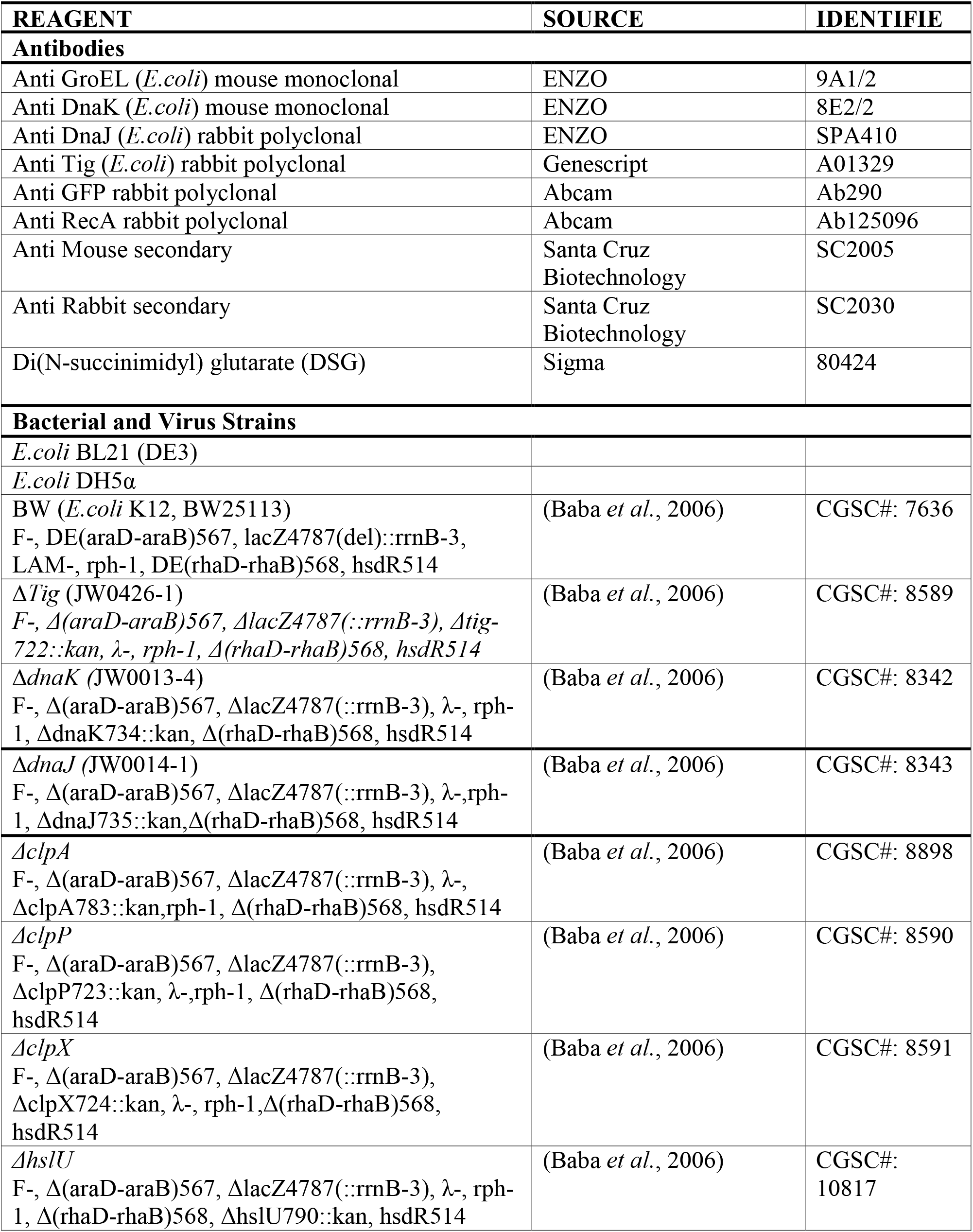
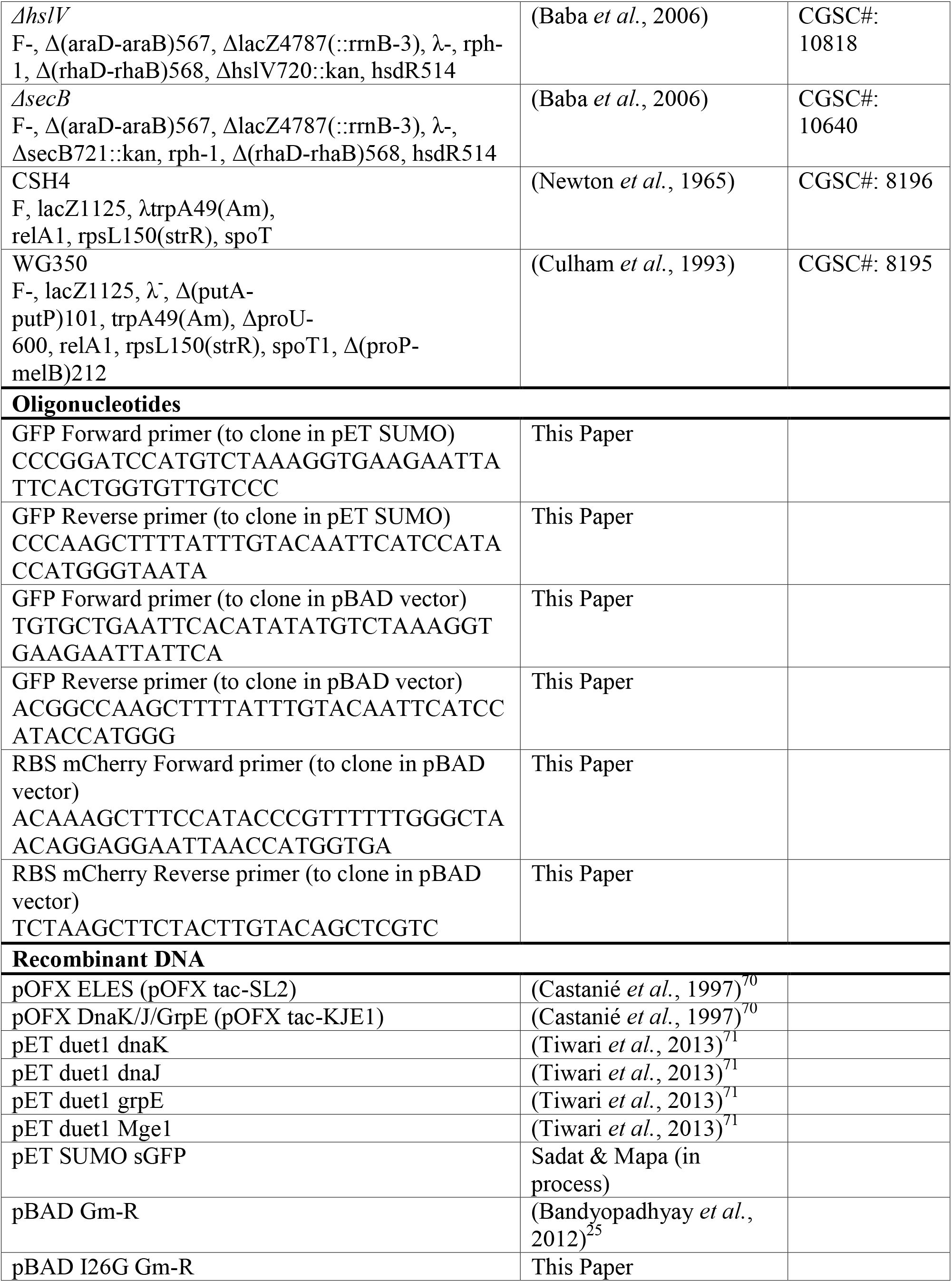
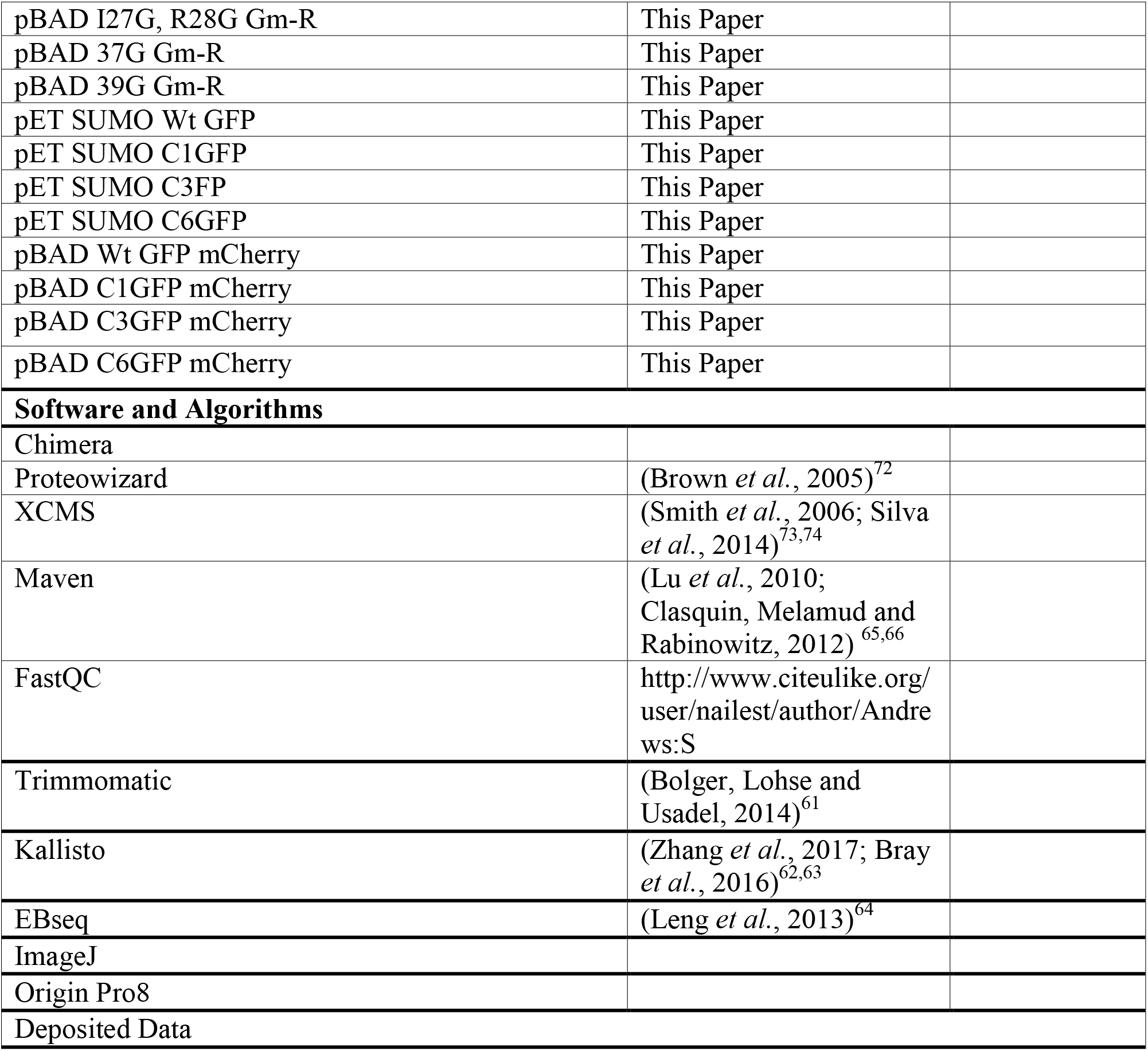

**FIGURE S1:**
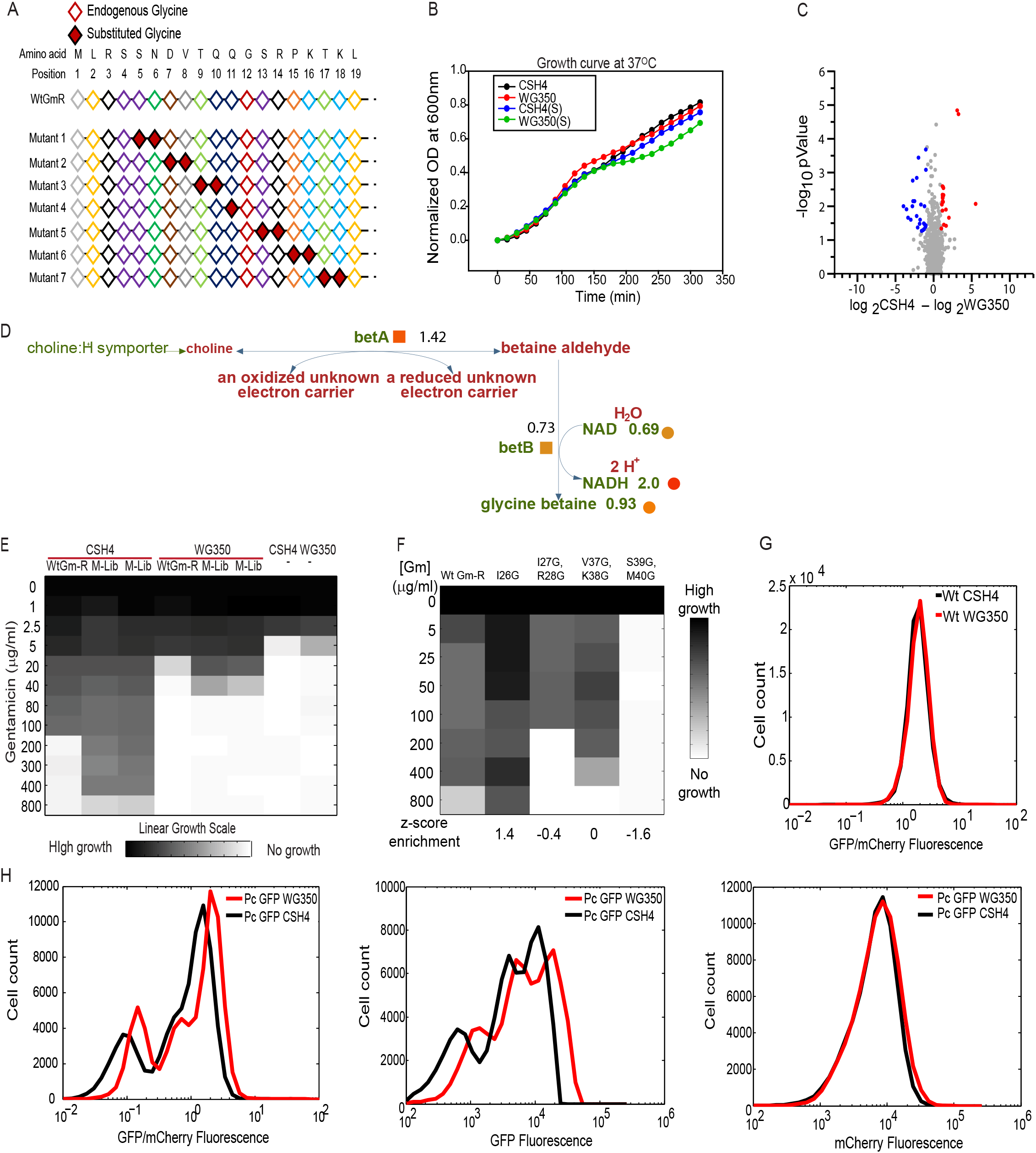
Strain specific metabolic differences lead to differential mutation buffering capacity (Related to Figure 2) **A.** Schematic of Glycine doublet substitution library (GG) of Gm-R mutants where two consecutive amino acids have been substituted with two Glycine residues (red solid filled) starting from 5th amino acid. **B.** Growth curve for *E.coli* strains CSH4 (black, blue) and WG350 (red, green) in presence and absence of 350 mM NaCl added in excess to the LB media while growing at 37°C. **C.** Scatter plot for log_2_ of fold change in metabolite concentrations between CSH4 and WG350 against −log_10_ of p-value. The significantly altered metabolites (p-value < 0.05, 5 biological replicates for each sample) are shown in colored circles D. Pathway for production of glycine betaine in *E.coli* showing correlation between metabolomics and transcriptomics. Metabolites upregulated are shown as circles and transcripts for the genes are shown as squares. Color intensity corresponds to fold upregulation **E.** Growth of untransformed CSH4 (-), WG350 (-) and cells transformed with Wt Gm-R and Glycine doublet mutant library plasmids (M-Lib) in increasing concentrations of Gentamicin (0-800 mg/ml). Increase in growth is shown as increase in color density. **F.** Correlation between NGS based quantification of enrichment scores (z-score) and MIC based semi-quantitative activity measurement for Wt Gm-R and mutants I26G, I27G-R28G, V37G-K38G, S39G-K40G of Glycine doublet mutant library. **G.** Histogram for *in vivo* fluorescence of Wt GFP (represented as ratio of GFP/mCherry) in CSH4 (black) and WG350 (red) **H.** Histogram for ratio of GFP/mCherry fluorescence (left) and independent fluorescence of GFP (middle) and mCherry channel (right) of mutant library Pc in CSH4 and WG350.

**FIGURE S2:**
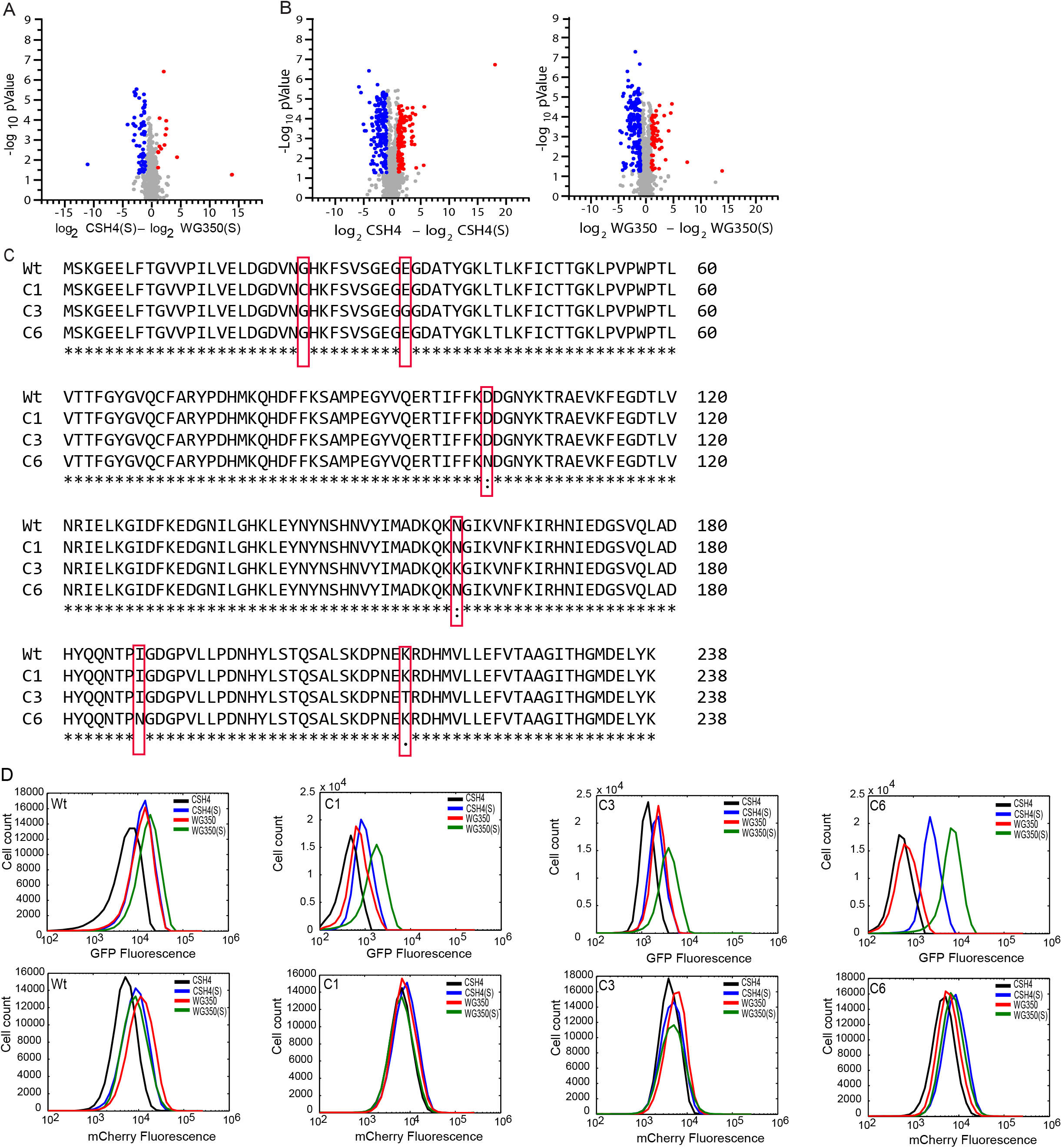
Osmotic stress induced modification in cellular metabolism changes the subset of mutations buffered (Related to Figure 3) **A.** Scatter plot for log_2_ of fold change in metabolic features between WG350 and CSH4 during osmotic shock against p-value. The significantly different metabolites (p-value < 0.05, 5 biological replicates for each sample) are represented in colored circles. **B.** Scatter plot for log_2_ of fold change in metabolic features within strains CSH4 (left) and WG350 (right) on osmotic shock(s) against p-value. The significantly different metabolites (p-value < 0.05, 5 biological replicates for each sample) are represented in colored circles. **C.** Amino acid sequence alignment of Wt GFP (YeGFP) with the isolated mutants C1, C3 and C6. Mutations are shown in red boxes. **D.**Histogram for GFP (top panel) and mCherry fluorescence (bottom panel) of the Isolated clones (Wt, C1, C3 and C6 GFP) in WG350 and CSH4 in the presence and absence of 350 mM NaCl.

**FIGURE S3:**
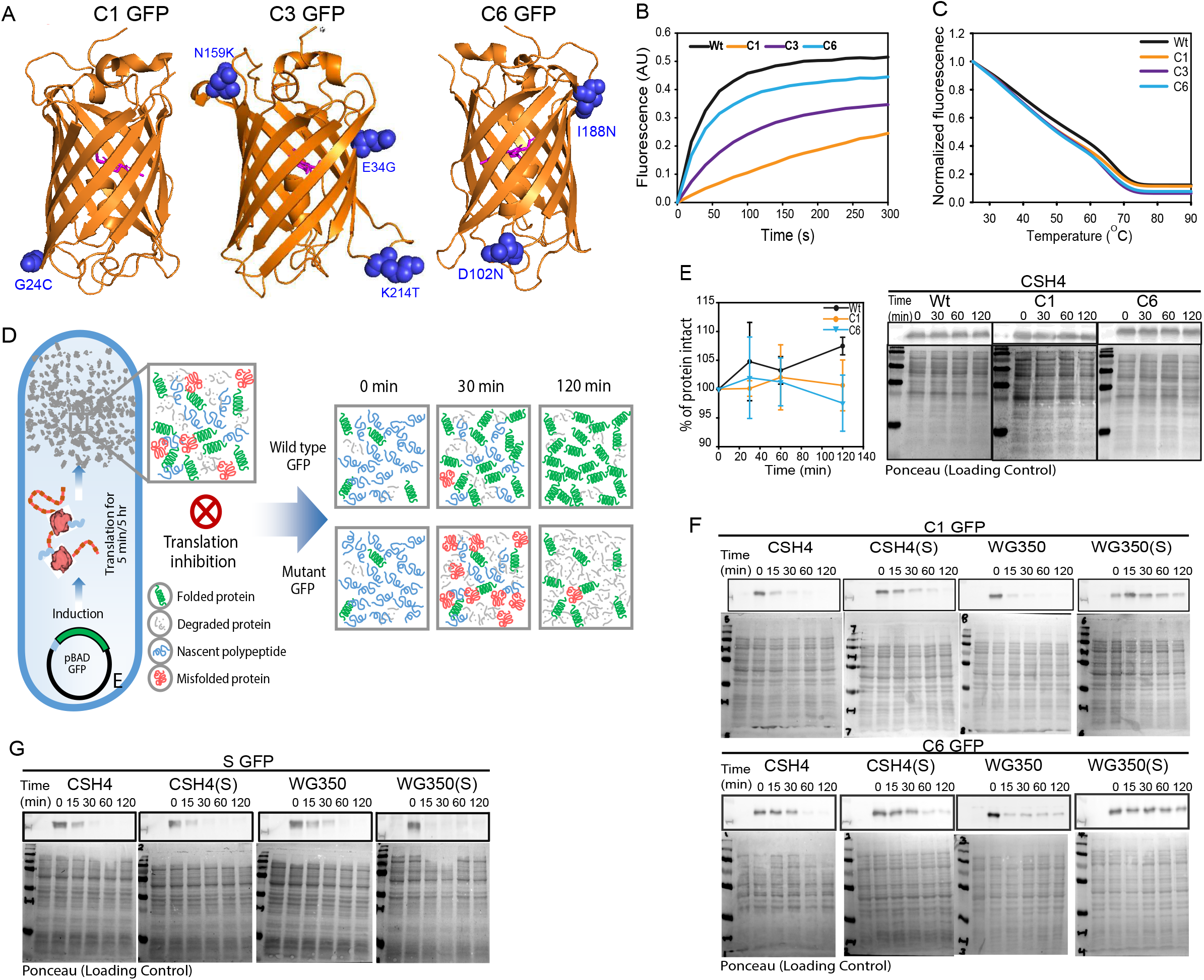
Fluorescence increase is not due better proteostasis in WG350(S) (Related to Figure 4) **A.** Mutations buffered in WG350 while growing in 350 mM NaCl (C1, C3, and C6) are marked on the GFP crystal structure (pdb: 1GFL) (Yang, Moss and Phillips, 1996). Fluorophore at the centre of the barrel is highlighted in magenta. **B.** Refolding traces of Wt and GFP mutants obtained on unfolding the proteins in 6 M GuHCl, followed by a 100-fold dilution into Buffer-A (50 mM Tris, 150 mM NaCl, 2 mM DTT, pH 7.4) to a final concentration of 200 nm for the proteins. The GFP fluorescence (AU) is plotted against time (s). **C.** Average thermal melt curves for Wt, C1, C3 and C6 GFP. Fluorescence of each mutants at different temperatures has been normalized to its respective fluorescence at 25°C. **D.** Schematic of Chloramphenicol-based chase for degradation of GFP as described in star methods. After a short (5 min) induction of GFP with 0.5% arabinose, translation was stalled with Chloramphenicol. Cells were lysed at different points and the amount of intact GFP was monitored by immunoblotting with anti-GFP antibody. **E. P**lot for the gel-based quantification of intact GFP (left) and representative images for immunoblotting to chase degradation of Wt, C1 and C6 GFP in CSH4 after 5 hr induction of GFP with 0.5% arabinose followed by translation arrest with Chloramphenicol. Ponceau has been used as loading control. **F.** Representative images for immunoblotting to chase degradation of nascent chains of C1 and C6 GFP as described in Star Methods in CSH4 and WG350 in presence and absence of osmotic stress. GFP degrades slowest in WG350(S). Ponceau has been used as loading control. **G.** Representative images for immunoblotting to chase degradation of sGFP in CSH4 and WG350 in presence and absence of osmotic stress. Ponceau has been used as loading control.

**Figure S4:**
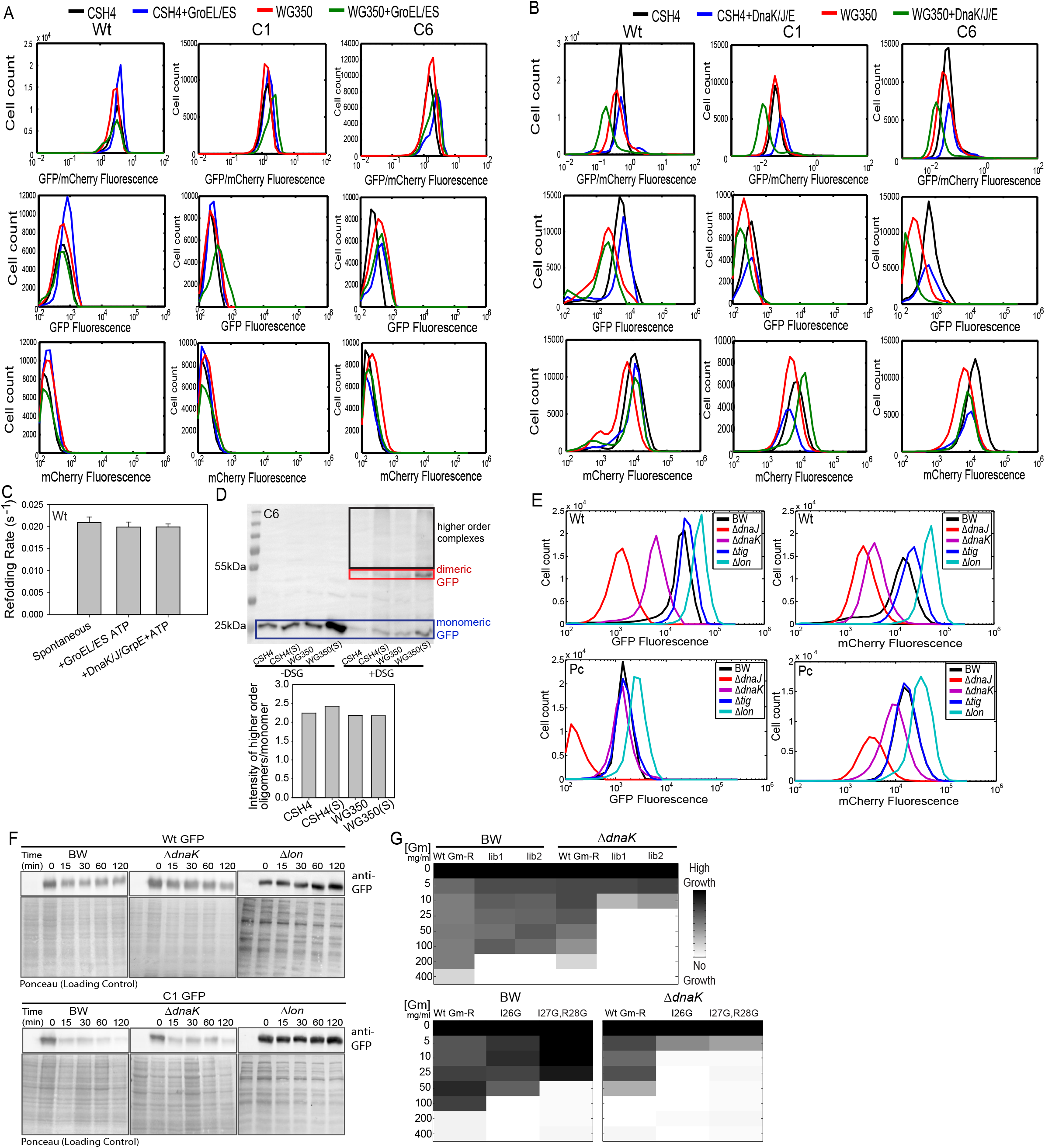
Observed buffering of GFP mutants is not channeled only through molecular chaperones (Related to Figure 5) **A.** Histogram for GFP/mCherry Fluorescence (top panel), GFP (middle panel) and mCherry fluorescence (bottom panel) in absence of overexpressed GroEL/ES (black, red, 2 biological replicates) and while co-expressed with GroEL/ES (blue, green, 2 biological replicates) in CSH4 and WG350. Induction protocol was followed as described in Star Methods. **B.** Histogram for GFP/mCherry Fluorescence (top panel), GFP (middle panel) and mCherry fluorescence (bottom panel) in absence of overexpressed DnaK/J/GrpE (black, red, 2 biological replicates) and while co-expressed with DnaK/J/GrpE (blue, green, 2 biological replicates) in CSH4 and WG350. **C.** Representative images for immunoblotting to chase degradation of Wt and C1 GFP in WT *E.coli* K12 strain BW25113 (BW) chaperone (Δ*dnaK*), and protease (Δ*lon*) knockout strain. Ponceau has been used as loading control. **D.** Immunoblotting for GFP showing the recruitment of nascently formed C6 GFP into multiprotein complexes. Free (or monomeric) GFP are shown with blue box, dimeric GFP are shown in red box and the multimeric complexes are shown in black box. The ratio of GFP in multimeric complex and the amount in monomeric state is shown in the accompanying bar graph. **E.** Histogram of GFP/mCherry fluorescence of Wt GFP and pool of GFP mutants having compromised fluorescence (Pc) in BW and molecular chaperone knockout strains (Δ*dnaJ*, Δ*dnaK*, Δ*tig*, Δ*lon*). **I.** Representative images for immunoblotting of chase degradation of Wt and C1 GFP in BW, chaperone (Δ*dnaK*), and protease (Δ*lon*) knockout strain. Ponceau has been used as loading control. **G.** Heatmap for growth based activity of Wt and Gm-R mutant library (lib1, lib2) (top panel), Wt Gm-R anf Gm-R mutants (I26G, I27G-R28G) in BW and molecular chaperone knockout strain (Δ*dnaK*) in increasing concentrations of gentamicin.

**Figure S5:**
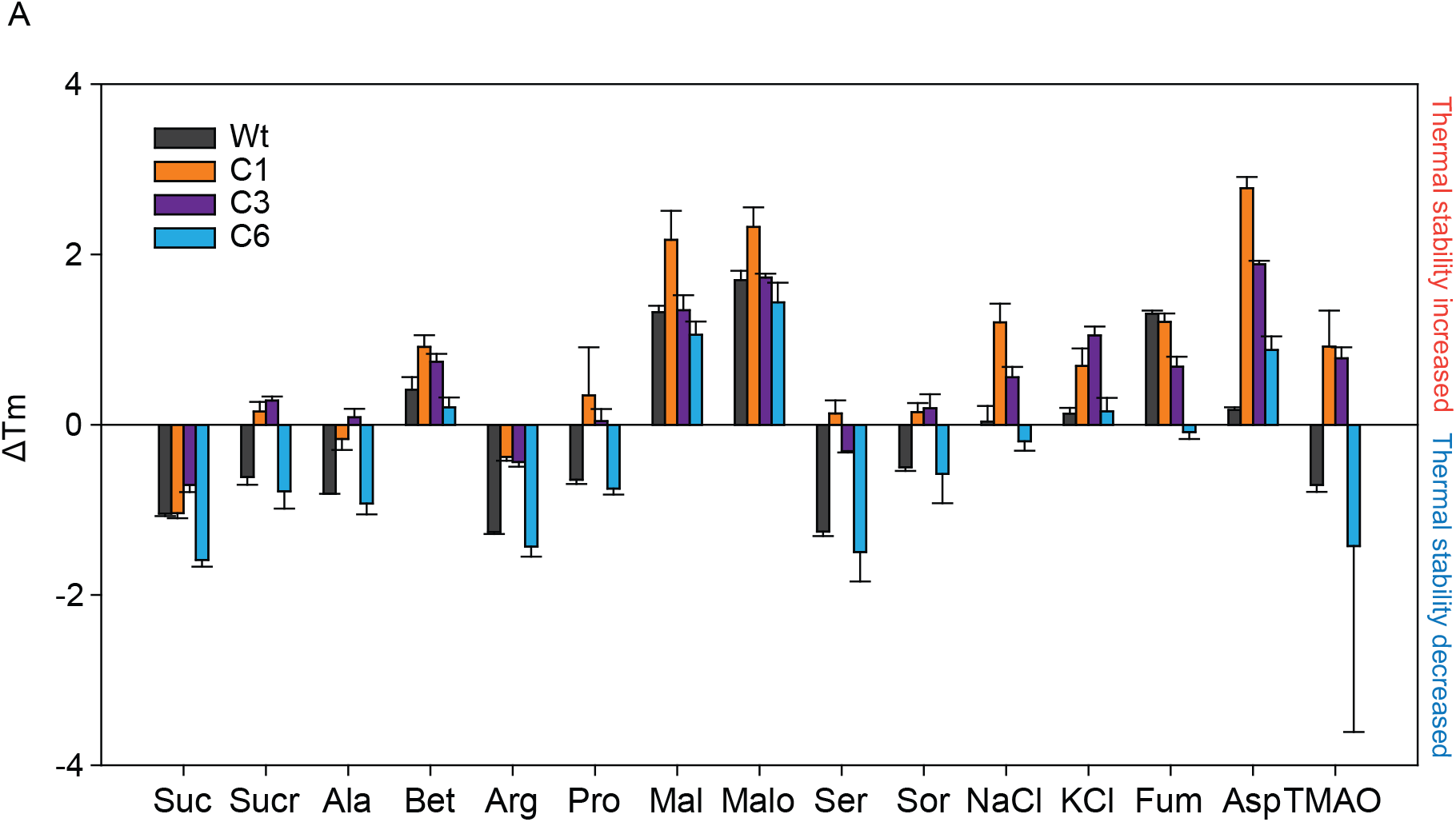
Different small molecules affect energetics of folding in a mutant specific manner (Related to Figure 6) **A.** Thermal melting profile of Wt, C1, C3 and C6 GFP as change in melting temperature (ΔTm) in presence of different small molecules. The change in Tm in presence of a small molecule is obtained by subtracting Tm in presence of Buffer-A (50 mM Tris, 150 mM NaCl, 2 mM DTT, pH 7.4) from Tm in presence of that respective small molecule (100 mM). Error bars indicate standard deviation among 3 technical replicates. Suc: Succinate, Sucr: Sucrose, Ala: Alanine, Bet: Betaine, Arg: Arginine, Pro: Proline, Mal: Malate, Malo: Malonate, Ser: Serine, Sor: Sorbitol, Fum: Fumarate, Asp: Aspartate, TMAO: Trimethylamine-N-oxide

**Figure S6:**
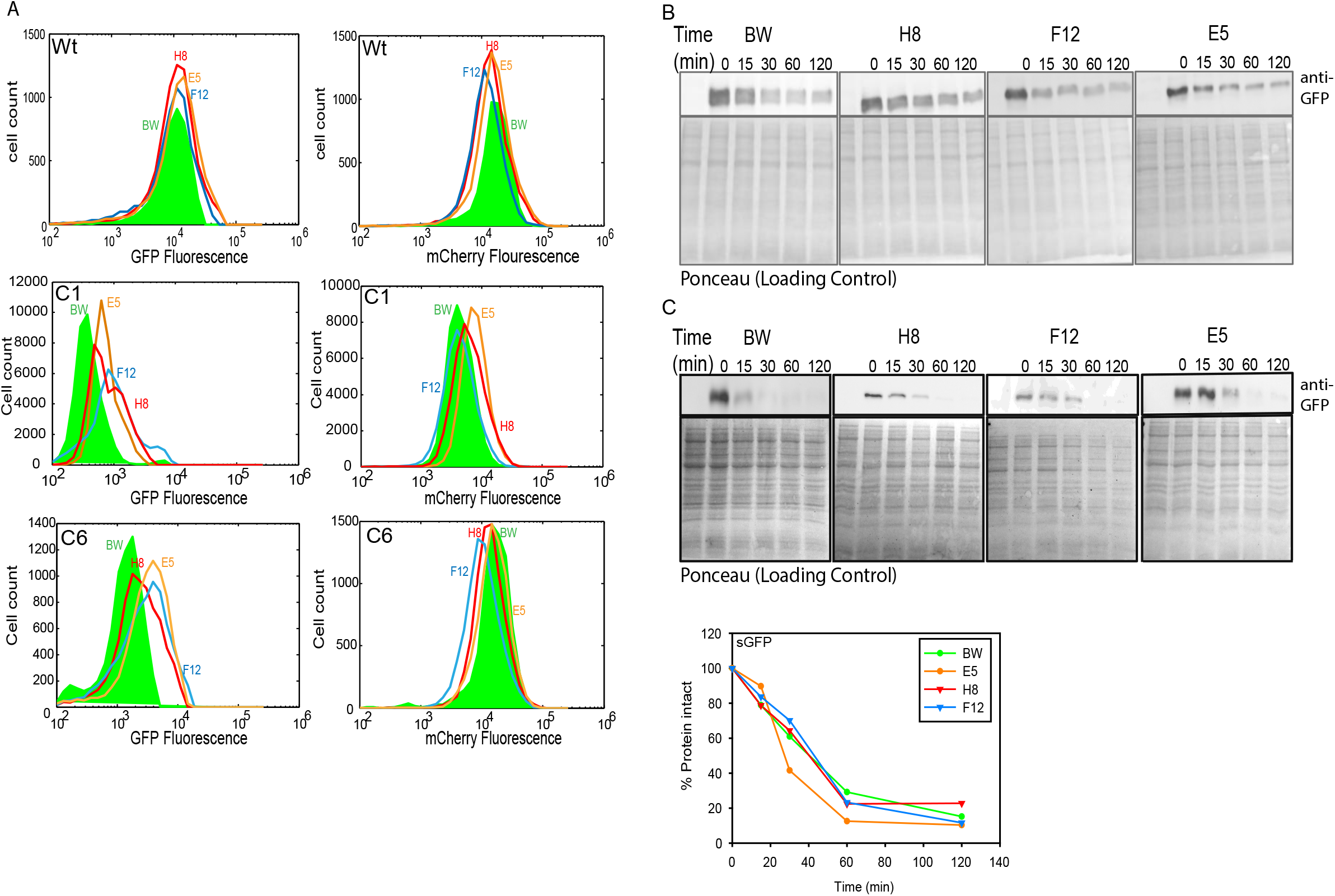
Modification in metabolome due to evolution changes the mutation buffering capacity of the cell (Related to Figure 8) **A.** Histogram for fluorescence of Wt and GFP mutants C1 and C6 represented as GFP (left panel) and mCherry fluorescence (right panel) in unevolved BW (WT E.coli K 12, BW25113) and evolved osmotolerant strains E5, F12, H8. **B.** Representative images of Chloramphenicol based chase for checking degradation rates of C1 GFP in BW (WT *E.coli* K 12, BW25113) and evolved strains E5, F12, H8. GFP is degraded slower in evolved strains than BW **C.** Representative images of Chloramphenicol based chase for sGFP in BW (WT *E.coli* K 12, BW25113) and evolved strains E5, F12, H8. sGFP is degraded with similar rates in evolved and unevolved BW as shown in densitometric analysis plot at the bottom

**Table S1:** List of metabolites along with their log_2_ of fold change and p-values obtained by untargeted metabolite profiling and analysis using MAVEN platform (Related to Figure 2 and 3)

| S.n<br>o. | med<br>Mz | med<br>Rt | Max<br>Quali<br>ty | compound | Log<sub>2</sub> fold change |  |  |  | Log<sub>10</sub> p-value |  |  |  |
| --- | --- | --- | --- | --- | --- | --- | --- | --- | --- | --- | --- | --- |
|  |  |  |  |  | CSH4<br>-<br>WG350 | CSH4<br>-<br>CSH4<br>(S) | WG350<br>-<br>WG350<br>(S) | CSH4<br>(S)-<br>WG350<br>(S) | CSH4-WG350 | CSH4-CSH4<br>(S) | WG350<br>-<br>WG350<br>(S) | CSH4<br>(S)-<br>WG350<br>(S) |
| 1 | 363.0344 | 2.041682 | 0.848391 | N-Acetylputrescine | 0.00 | -14.03 | -14.29 | -0.26 | 0.01 | 0.97 | 1.54 | 0.45 |
| 2 | 283.0684 | 4.029608 | 0.849596 | Geranyl-PP | 0.00 | -13.99 | -13.89 | 0.10 | 0.05 | 1.67 | 1.43 | 0.20 |
| 3 | 151.025 | 1.858412 | 0.819399 | fructose-1-6-bisphosphate | 0.00 | -12.65 | 0.00 | 12.65 | 0.00 | 2.24 | 0.43 | 5.16 |
| 4 | 116.0705 | 2.341379 | 0.828605 | succinate | 0.00 | -11.67 | -25.78 | -14.11 | 0.43 | 1.60 | 2.15 | 1.80 |
| 5 | 243.0622 | 2.131804 | 0.848934 | UMP | 2.10 | -4.63 | -4.86 | 1.87 | 0.49 | 2.56 | 1.80 | 1.80 |
| 6 | 167.0204 | 2.374183 | 0.847362 | orotidine-5-phosphate | 10.98 | -4.25 | -14.62 | 0.61 | 1.12 | 4.78 | 1.71 | 1.38 |
| 7 | 323.0291 | 3.537884 | 0.851401 | betaine | -0.93 | -4.17 | -7.94 | -4.70 | 0.53 | 0.95 | 0.98 | 1.78 |
| 8 | 606.0748 | 4.036123 | 0.851434 | Octoluse Bisphosphate | 1.06 | -3.88 | -4.69 | 0.24 | 0.15 | 1.00 | 1.04 | 0.28 |
| 9 | 565.0482 | 4.056136 | 0.857477 | tryptophan | 0.44 | -3.75 | -4.18 | 0.01 | 0.05 | 1.93 | 1.71 | 0.22 |
| 10 | 402.9949 | 2.973083 | 0.84958 | thiamine-phosphate | 0.12 | -3.73 | -2.99 | 0.87 | 0.01 | 1.34 | 1.15 | 3.48 |
| 11 | 180.0657 | 2.219842 | 0.858241 | fumarate | -2.93 | -3.27 | -6.82 | -6.49 | 0.88 | 0.51 | 3.44 | 1.03 |
| 12 | 203.0819 | 1.413762 | 0.840261 | FMN | 1.54 | -2.86 | -1.06 | 3.34 | 1.17 | 1.98 | 0.65 | 4.25 |
| 13 | 203.0819 | 1.505575 | 0.858171 | asparagine | -1.04 | -2.73 | 0.10 | 1.79 | 0.62 | 2.12 | 0.10 | 3.83 |
| 14 | 341.1089 | 2.038149 | 0.846411 | glutathione disulfide | -1.96 | -2.52 | -0.13 | 0.43 | 1.29 | 1.15 | 0.39 | 0.24 |
| 15 | 421.0756 | 1.887725 | 0.851835 | CMP | -1.08 | -2.37 | 1.97 | 3.26 | 0.47 | 1.60 | 1.10 | 1.82 |
| 16 | 241.0827 | 2.019653 | 0.856852 | GMP | 1.22 | -2.33 | -2.11 | 1.44 | 1.97 | 4.11 | 2.86 | 3.62 |
| 17 | 344.0691 | 2.006949 | 0.846896 | UDP | -0.13 | -2.28 | -1.68 | 0.47 | 0.10 | 4.33 | 3.04 | 0.73 |
| 18 | 424.039 | 1.499295 | 0.860667 | aspartate | -0.40 | -1.99 | 2.16 | 3.75 | 0.64 | 3.32 | 2.82 | 5.07 |
| 19 | 264.1045 | 1.446013 | 0.858174 | trehalose/sucrose | 1.55 | -1.89 | -1.60 | 1.84 | 3.11 | 3.67 | 2.71 | 3.08 |
| 20 | 866.<br>124<br>9 | 1.450<br>47 | 0.857<br>708 | Cellobiose | 1.55 | -1.89 | -1.60 | 1.84 | 3.11 | 3.67 | 2.71 | 3.08 |
| 21 | 117.<br>018<br>6 | 2.093<br>047 | 0.854<br>168 | N-Acetyl-L-alanine | -1.80 | -1.77 | 0.08 | 0.05 | 0.78 | 1.01 | 0.38 | 0.06 |
| 22 | 171.<br>006<br>5 | 3.843<br>684 | 0.849<br>186 | IMP | -0.41 | -1.66 | -1.07 | 0.18 | 0.38 | 2.09 | 1.54 | 0.45 |
| 23 | 253.<br>009<br>9 | 2.067<br>247 | 0.861<br>783 | Pyrogluta<br>mic acid | -1.85 | -1.53 | -0.12 | -0.43 | 0.64 | 0.67 | 0.41 | 0.80 |
| 24 | 368.<br>998<br>9 | 3.985<br>863 | 0.849<br>202 | S-methyl-5--<br>thioadenos<br>ine | -0.27 | -1.28 | -1.28 | -0.28 | 0.15 | 1.34 | 1.39 | 0.43 |
| 25 | 266.<br>070<br>7 | 1.685<br>908 | 0.833<br>351 | ribose-<br>phosphate | -0.60 | -1.15 | 0.82 | 1.37 | 0.75 | 1.65 | 1.10 | 2.80 |
| 26 | 296.<br>080<br>6 | 2.066<br>397 | 0.852<br>619 | N-acetyl-L-ornithine | 0.30 | -1.10 | -1.11 | 0.30 | 0.35 | 2.75 | 2.67 | 0.64 |
| 27 | 383.<br>115<br>2 | 3.042<br>196 | 0.771<br>733 | Acetyllysin<br>e | 0.05 | -0.96 | -0.98 | 0.04 | 0.11 | 1.80 | 2.12 | 0.13 |
| 28 | 229.<br>011<br>2 | 2.020<br>989 | 0.853<br>25 | adenosine | 0.89 | -0.91 | -1.32 | 0.48 | 0.93 | 2.14 | 2.15 | 2.26 |
| 29 | 375.<br>131 | 2.323<br>842 | 0.847<br>81 | xanthosine<br>-5-phosphate | -0.18 | -0.89 | -0.73 | -0.01 | 0.04 | 1.82 | 2.08 | 0.03 |
| 30 | 176.<br>935 | 4.759<br>889 | 0.852<br>717 | 2-Isopropylm<br>alic acid | 0.23 | -0.74 | -0.66 | 0.30 | 0.24 | 1.32 | 1.66 | 0.68 |
| 31 | 128.<br>034<br>1 | 1.833<br>35 | 0.853<br>449 | dCDP | -0.36 | -0.47 | 0.46 | 0.58 | 0.07 | 1.29 | 0.49 | 2.60 |
| 32 | 168.<br>065<br>6 | 2.160<br>493 | 0.853<br>901 | xanthosine | 0.45 | -0.45 | -0.52 | 0.37 | 1.07 | 1.80 | 1.32 | 1.07 |
| 33 | 114.<br>055<br>1 | 10.95<br>572 | 0.731<br>41 | p-<br>hydroxybe<br>nzoate | -0.33 | -0.42 | -0.22 | -0.14 | 0.67 | 0.25 | 1.34 | 0.49 |
| 34 | 225.<br>039<br>2 | 11.50<br>664 | 0.848<br>89 | Sedohepto<br>luse<br>bisphosph<br>ate | -0.26 | -0.24 | -0.07 | -0.09 | 0.28 | 0.00 | 0.14 | 0.08 |
| 35 | 145.<br>030<br>1 | 1.693<br>76 | 0.833<br>608 | cystathioni<br>ne | 0.19 | -0.23 | -1.06 | -0.64 | 0.54 | 0.38 | 1.94 | 0.55 |
| 36 | 218.<br>102<br>9 | 1.981<br>234 | 0.860<br>063 | hypoxanthi<br>ne | -0.22 | -0.19 | -0.01 | -0.04 | 0.52 | 0.74 | 0.11 | 0.03 |
| 37 | 137.<br>023<br>2 | 1.572<br>158 | 0.857<br>927 | glutamine | -0.07 | -0.16 | -0.09 | 0.00 | 0.28 | 0.01 | 0.07 | 0.06 |
| 38 | 367.<br>019<br>8 | 1.393<br>888 | 0.848<br>99 | UDP-D-glucose | -0.14 | -0.14 | -0.10 | -0.09 | 0.64 | 0.59 | 0.14 | 0.02 |
| 39 | 131.<br>081<br>4 | 6.987<br>23 | 0.856<br>877 | Kynurenin<br>e | 0.22 | -0.11 | -0.34 | -0.02 | 0.22 | 0.36 | 1.13 | 0.16 |
| 40 | 399.<br>013<br>1 | 5.573<br>004 | 0.850<br>927 | riboflavin | -0.10 | -0.09 | -0.13 | -0.14 | 0.07 | 0.04 | 0.50 | 0.29 |
| 41 | 319.<br>046<br>3 | 5.199<br>954 | 0.849<br>735 | Thiamine<br>pyrophosp<br>hate | -0.03 | -0.09 | -0.05 | 0.01 | 0.12 | 0.54 | 0.56 | 0.12 |
| 42 | 146.<br>044<br>7 | 5.076<br>94 | 0.840<br>613 | Octoluse<br>8/1P | 0.10 | -0.08 | -0.32 | -0.14 | 0.20 | 0.50 | 0.53 | 0.25 |
| 43 | 742.<br>069<br>5 | 3.301<br>201 | 0.735<br>053 | valine | 0.22 | -0.08 | -0.36 | -0.06 | 0.27 | 0.36 | 1.11 | 0.22 |
| 44 | 664.<br>118 | 1.939<br>427 | 0.662<br>212 | cyclic-AMP | 0.27 | -0.02 | 0.62 | 0.91 | 0.55 | 0.64 | 1.54 | 1.10 |
| 45 | 662.<br>102<br>7 | 8.182<br>36 | 0.852<br>802 | tryptophan | 0.28 | -0.01 | -0.40 | -0.11 | 0.20 | 0.13 | 0.99 | 0.61 |
| 46 | 175.<br>035<br>1 | 1.743<br>987 | 0.851<br>61 | glucose-6-phosphate | 0.35 | 0.04 | 1.96 | 2.27 | 0.75 | 0.07 | 0.43 | 3.75 |
| 47 | 129.<br>102<br>3 | 6.072<br>19 | 0.850<br>308 | thiamine | 0.00 | 0.05 | -0.08 | -0.14 | 0.13 | 0.05 | 0.50 | 0.34 |
| 48 | 173.<br>092<br>2 | 2.708<br>844 | 0.814<br>59 | pantothena<br>te | 0.40 | 0.06 | -0.53 | -0.18 | 0.33 | 0.03 | 1.38 | 0.97 |
| 49 | 130.<br>049<br>8 | 5.998<br>872 | 0.854<br>558 | biotin | 0.14 | 0.06 | -0.17 | -0.09 | 0.19 | 0.08 | 0.77 | 0.16 |
| 50 | 187.<br>071<br>6 | 1.777<br>959 | 0.832<br>413 | shikimate-3-phosphate | 0.04 | 0.06 | 0.20 | 0.18 | 0.44 | 0.47 | 1.26 | 0.37 |
| 51 | 188.<br>055<br>6 | 1.504<br>867 | 0.772<br>704 | N-acetyl-<br>glutamine | 0.04 | 0.09 | 0.16 | 0.10 | 0.02 | 0.01 | 1.50 | 1.00 |
| 52 | 300.<br>046<br>6 | 1.719<br>046 | 0.848<br>407 | guanine | -0.13 | 0.13 | 0.24 | -0.02 | 0.45 | 0.03 | 0.79 | 0.22 |
| 53 | 179.<br>055<br>2 | 1.501<br>844 | 0.851<br>44 | inosine | -0.21 | 0.17 | 1.76 | 1.37 | 0.00 | 0.07 | 2.49 | 1.18 |
| 54 | 134.<br>029<br>4 | 2.005<br>65 | 0.855<br>251 | dGMP | 0.40 | 0.25 | 0.12 | 0.27 | 0.68 | 0.33 | 0.71 | 0.24 |
| 55 | 148.<br>042<br>6 | 3.287<br>45 | 0.850<br>251 | D-sedoheptul<br>ose-1/7-phosphate | 0.19 | 0.26 | 0.12 | 0.05 | 0.03 | 0.17 | 0.99 | 0.45 |
| 56 | 133.<br>013<br>1 | 1.936<br>306 | 0.677<br>158 | methionine | -0.17 | 0.29 | 0.50 | 0.05 | 0.16 | 1.54 | 0.85 | 0.13 |
| 57 | 145.<br>097<br>1 | 1.390<br>545 | 0.850<br>053 | UDP-N-acetyl-<br>glucosami<br>ne | -0.21 | 0.40 | 0.18 | -0.42 | 0.17 | 0.40 | 0.66 | 0.17 |
| 58 | 205.<br>034<br>7 | 2.536<br>442 | 0.851<br>966 | proline | -0.88 | 0.40 | -1.20 | -2.48 | 0.33 | 0.24 | 0.81 | 1.84 |
| 59 | 289.<br>115<br>5 | 1.833<br>144 | 0.851<br>898 | a-<br>ketoglutar<br>ate | 0.38 | 0.42 | 1.06 | 1.02 | 1.23 | 1.41 | 2.15 | 1.78 |
| 60 | 207.<br>077 | 2.012<br>048 | 0.833<br>829 | tyrosine | 0.15 | 0.48 | 0.51 | 0.18 | 0.70 | 2.96 | 2.09 | 1.13 |
| 61 | 267.<br>072<br>5 | 1.519<br>962 | 0.766<br>428 | Aminoadipi<br>c acid | 0.22 | 0.49 | 0.01 | -0.26 | 0.38 | 1.51 | 0.22 | 1.52 |
| 62 | 347.<br>039<br>5 | 2.047<br>51 | 0.799<br>432 | N-acetyl-<br>glutamate | 0.26 | 0.53 | 0.47 | 0.19 | 0.21 | 1.07 | 0.92 | 0.36 |
| 63 | 135.<br>030<br>1 | 3.948<br>531 | 0.850<br>648 | acetyl-CoA | 0.12 | 0.55 | 1.77 | 1.34 | 0.04 | 2.12 | 2.55 | 2.32 |
| 64 | 112.<br>428<br>6 | 1.409<br>344 | 0.858<br>314 | arginine | 0.12 | 0.56 | 0.53 | 0.09 | 0.51 | 1.82 | 2.03 | 0.09 |
| 65 | 154.<br>061<br>1 | 1.914<br>433 | 0.856<br>739 | glutathione | 0.05 | 0.57 | 0.97 | 0.45 | 0.03 | 1.40 | 2.23 | 1.89 |
| 66 | 282.<br>086<br>1 | 1.459<br>852 | 0.857<br>308 | ornithine | -0.04 | 0.58 | 0.55 | -0.07 | 0.34 | 0.41 | 2.33 | 0.09 |
| 67 | 150.<br>041<br>1 | 2.037<br>107 | 0.853<br>763 | thymidine | 0.18 | 0.59 | 0.36 | -0.04 | 0.44 | 1.94 | 2.04 | 0.31 |
| 8 | 362.<br>051<br>1 | 1.499<br>073 | 0.851<br>382 | L-arginino-<br>succinate | -0.38 | 0.63 | 2.28 | 1.27 | 0.38 | 1.49 | 2.82 | 0.84 |
| 69 | 611.<br>140<br>8 | 2.658<br>341 | 0.853<br>422 | N-acetyl-<br>glucosami-<br>ne-1/6-phosphate | 0.02 | 0.67 | 0.51 | -0.14 | 0.21 | 2.95 | 2.74 | 0.61 |
| 70 | 306.<br>076<br>5 | 1.728<br>165 | 0.823<br>201 | N-carbamoyl-L-aspartate | 0.61 | 0.73 | 0.23 | 0.11 | 0.87 | 0.40 | 0.17 | 0.02 |
| 71 | 145.<br>060<br>7 | 2.068<br>78 | 0.715<br>853 | uridine | 0.12 | 0.75 | 0.56 | -0.06 | 0.21 | 2.06 | 1.65 | 0.42 |
| 72 | 259.<br>022 | 1.509<br>566 | 0.783<br>648 | pyridoxine | -0.08 | 0.76 | 0.58 | -0.26 | 0.11 | 3.64 | 3.39 | 2.73 |
| 73 | 178.<br>071<br>4 | 1.853<br>829 | 0.854<br>485 | sn-<br>glycerol-3-phosphate | 0.08 | 0.83 | 0.36 | -0.39 | 0.00 | 2.49 | 1.50 | 1.29 |
| 74 | 177.<br>039<br>5 | 2.047<br>71 | 0.750<br>985 | xanthine | -0.02 | 0.84 | 0.62 | -0.24 | 0.23 | 2.54 | 2.09 | 1.04 |
| 75 | 313.<br>058<br>9 | 2.024<br>257 | 0.820<br>586 | citrate/isoci-<br>trate | 0.48 | 0.85 | 0.79 | 0.42 | 0.68 | 1.42 | 1.53 | 1.24 |
| 76 | 109.<br>526<br>4 | 1.386<br>311 | 0.855<br>701 | adenosine<br>5--<br>phosphosu-<br>lfate | 0.31 | 0.85 | 0.50 | -0.04 | 0.53 | 2.48 | 2.30 | 0.33 |
| 77 | 338.<br>989 | 1.485<br>925 | 0.818<br>313 | O-acetyl-L-serine | -0.14 | 0.93 | 0.85 | -0.22 | 0.73 | 3.92 | 3.35 | 1.01 |
| 78 | 455.<br>101<br>4 | 6.339<br>029 | 0.859<br>511 | 2_3-dihydroxyb-<br>enzoic acid | -1.45 | 0.94 | -0.60 | -2.99 | 2.03 | 1.58 | 1.11 | 3.67 |
| 79 | 784.<br>150<br>8 | 1.505<br>536 | 0.850<br>999 | cytidine | -0.05 | 0.96 | 0.86 | -0.15 | 0.01 | 4.59 | 3.03 | 0.30 |
| 80 | 321.<br>049<br>6 | 1.780<br>281 | 0.860<br>946 | malate | -0.10 | 0.99 | 0.15 | -0.95 | 0.12 | 1.90 | 0.05 | 4.07 |
| 81 | 346.<br>055<br>8 | 1.441<br>355 | 0.856<br>271 | lysine | 1.42 | 1.02 | -0.06 | 0.34 | 1.83 | 1.76 | 0.14 | 1.05 |
| 82 | 686.<br>142<br>3 | 1.567<br>711 | 0.853<br>065 | lipoate | 0.04 | 1.08 | 0.61 | -0.43 | 0.38 | 3.09 | 2.51 | 2.90 |
| 83 | 251.<br>077<br>6 | 4.043<br>802 | 0.849<br>58 | FAD | -0.14 | 1.11 | 0.56 | -0.69 | 0.23 | 3.17 | 0.87 | 1.06 |
| 84 | 250.<br>093<br>7 | 2.100<br>759 | 0.846<br>352 | NADP+ | 0.30 | 1.11 | 1.08 | 0.27 | 0.04 | 0.65 | 1.58 | 0.25 |
| 85 | 386.<br>017<br>2 | 1.486<br>119 | 0.856<br>649 | D-gluconate | -0.02 | 1.15 | 1.23 | 0.05 | 0.30 | 2.63 | 3.13 | 0.20 |
| 86 | 289.<br>035<br>2 | 1.507<br>047 | 0.851<br>621 | 2-dehydro-D-gluconate | -0.21 | 1.28 | 1.75 | 0.27 | 0.04 | 2.48 | 1.42 | 0.27 |
| 87 | 258.0375 | 1.453563 | 0.828056 | histidine | 0.30 | 1.30 | 1.00 | 0.01 | 0.46 | 2.73 | 3.55 | 0.11 |
| 88 | 195.0502 | 2.024109 | 0.852796 | dTMP | -0.94 | 1.31 | 1.59 | -0.66 | 1.19 | 1.38 | 2.20 | 1.03 |
| 89 | 199.0007 | 1.550444 | 0.856554 | homoserine | -2.41 | 1.32 | -1.46 | -5.18 | 0.76 | 0.25 | 0.97 | 1.45 |
| 90 | 242.078 | 2.027355 | 0.788688 | glucono-δ-lactone | 0.47 | 1.33 | 1.05 | 0.18 | 0.65 | 2.37 | 1.75 | 0.20 |
| 91 | 239.0147 | 1.507822 | 0.829343 | myo-inositol | -0.17 | 1.38 | 1.11 | -0.44 | 0.91 | 4.50 | 3.63 | 1.23 |
| 92 | 221.0598 | 1.489268 | 0.854986 | citrulline | -0.20 | 1.46 | 1.22 | -0.43 | 0.35 | 3.66 | 2.77 | 1.07 |
| 93 | 328.0454 | 3.954113 | 0.838432 | succinyl-CoA/methylmalonyl-CoA | -0.26 | 1.66 | 1.51 | -0.42 | 0.34 | 1.03 | 2.36 | 0.81 |
| 94 | 689.0877 | 2.038819 | 0.852447 | cyclic bis (3→5-) dimeric GMP | -0.18 | 1.68 | 1.61 | -0.24 | 2.83 | 3.02 | 2.29 | 0.14 |
| 95 | 766.1091 | 1.525817 | 0.835367 | Phenylpropionic acid | 0.14 | 1.75 | 1.23 | -0.37 | 0.00 | 0.58 | 1.85 | 0.47 |
| 96 | 322.0445 | 3.260484 | 0.848054 | coenzyme A | -1.70 | 1.77 | 1.48 | -1.99 | 1.48 | 2.14 | 1.76 | 3.87 |
| 97 | 174.0874 | 1.793035 | 0.851756 | D-erythrose-4-phosphate | 0.19 | 1.89 | 1.83 | 0.14 | 0.42 | 1.66 | 1.94 | 0.01 |
| 98 | 191.019 | 1.979981 | 0.851608 | Methylcysteine | 0.95 | 2.07 | 1.70 | 0.58 | 0.58 | 1.09 | 0.71 | 0.54 |
| 99 | 341.1089 | 1.499742 | 0.852927 | S-ribosyl-L-homocysteine-nega | 0.12 | 2.14 | 0.40 | -1.63 | 0.11 | 2.81 | 0.72 | 1.16 |
| 100 | 243.0804 | 1.504422 | 0.852072 | deoxyinosine | 0.31 | 2.21 | 1.33 | -0.57 | 0.79 | 2.89 | 2.21 | 0.13 |
| 101 | 116.0711 | 1.985581 | 0.85367 | NADH | -2.59 | 2.54 | 3.62 | -1.52 | 1.44 | 0.68 | 3.51 | 0.63 |
| 102 | 132.0291 | 1.759809 | 0.850709 | trehalose-6-Phosphate | 2.73 | 2.56 | 0.30 | 0.47 | 3.62 | 3.26 | 0.95 | 2.36 |
| 103 | 131.045 | 1.46988 | 0.853972 | deoxyadenosine | 0.33 | 2.69 | 1.93 | -0.43 | 0.47 | 4.44 | 2.27 | 0.21 |
| 104 | 173.1035 | 1.516454 | 0.850518 | D-glucosamine-6-phosphate | 0.68 | 2.77 | 1.61 | -0.49 | 0.69 | 1.86 | 0.67 | 0.30 |
| 105 | 160.0605 | 1.466613 | 0.852018 | glucosamine | 0.42 | 2.83 | 2.92 | 0.51 | 0.89 | 3.49 | 2.96 | 0.18 |
| 106 | 426.0121 | 2.080937 | 0.848104 | NAD+ | -0.69 | 3.58 | 2.11 | -2.15 | 0.07 | 2.46 | 0.79 | 1.78 |
| 107 | 266.091 | 1.819532 | 0.852153 | Pyrophosphate | 0.22 | 4.02 | 3.37 | -0.43 | 0.55 | 5.90 | 4.13 | 0.82 |
| 108 | 187.108 | 1.889188 | 0.850813 | Uric acid | -0.31 | 4.32 | 3.55 | -1.08 | 0.17 | 4.38 | 2.87 | 0.74 |
| 109 | 808.1196 | 2.336574 | 0.830013 | dephospho-CoA | -1.36 | 11.89 | 2.10 | -11.15 | 1.16 | 1.83 | 1.11 | 0.44 |
| 110 | 145.<br>013<br>2 | 2.367<br>599 | 0.838<br>086 | Cystine | 1.04 | 12.24 | 11.20 | 0.00 | 2.09 | 7.53 | 1.86 | 0.43 |
| 111 | 153.<br>018<br>2 | 1.509<br>54 | 0.803<br>857 | S-adenosyl<br>L-homoCyst<br>eine | -0.86 | 12.98 | 13.85 | 0.00 | 1.67 | 8.01 | 6.35 | #DIV/0! |
| 112 | 175.<br>060<br>2 | 1.598<br>831 | 0.850<br>469 | prephenat<br>e | 0.46 | 13.61 | 13.14 | 0.00 | 0.07 | 5.34 | 1.52 | 0.75 |
| 113 | 193.<br>034<br>6 | 1.511<br>09 | 0.831<br>777 | guanosine | 0.08 | 14.42 | 1.34 | -13.00 | 0.45 | 1.36 | 2.89 | 1.14 |

**Key:** med Mz: Median $m/z$ for metabolites, medRt: Median Retention Time, Max Quality: quality of peaks detected, compound: Metabolite, CSH4 - WG350: $\log_2$ of fold change and between CSH4 and WG350, CSH4 - CSH4 (S): $\log_2$ of fold change between CSH4 and CSH4 salt, WG350 - WG350 (S): $\log_2$ of fold change between WG350 and WG350 salt, CSH4 (S)- WG350 (S): $\log_2$ of fold change between CSH4 salt and WG350 salt

**Table S2:**
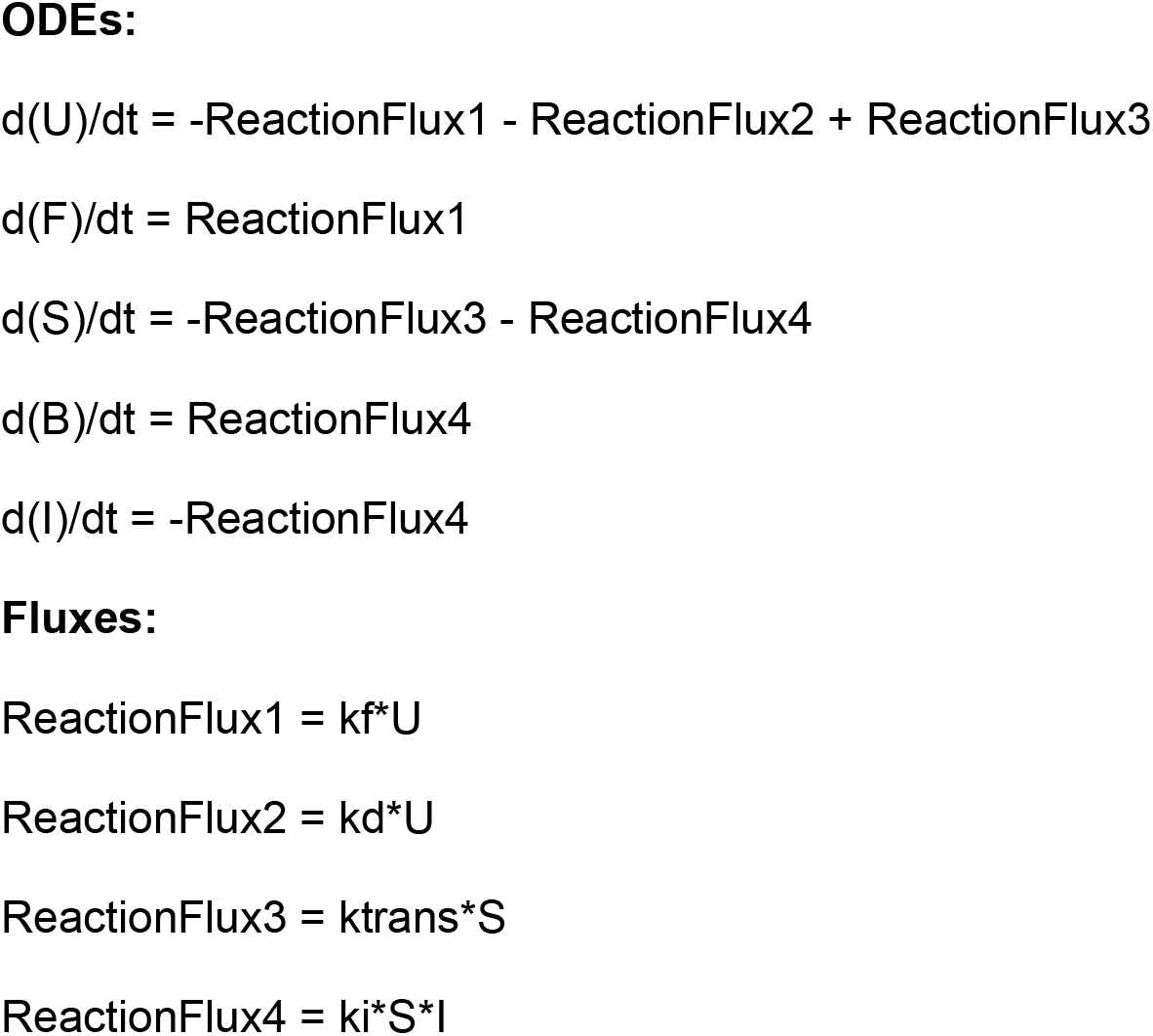
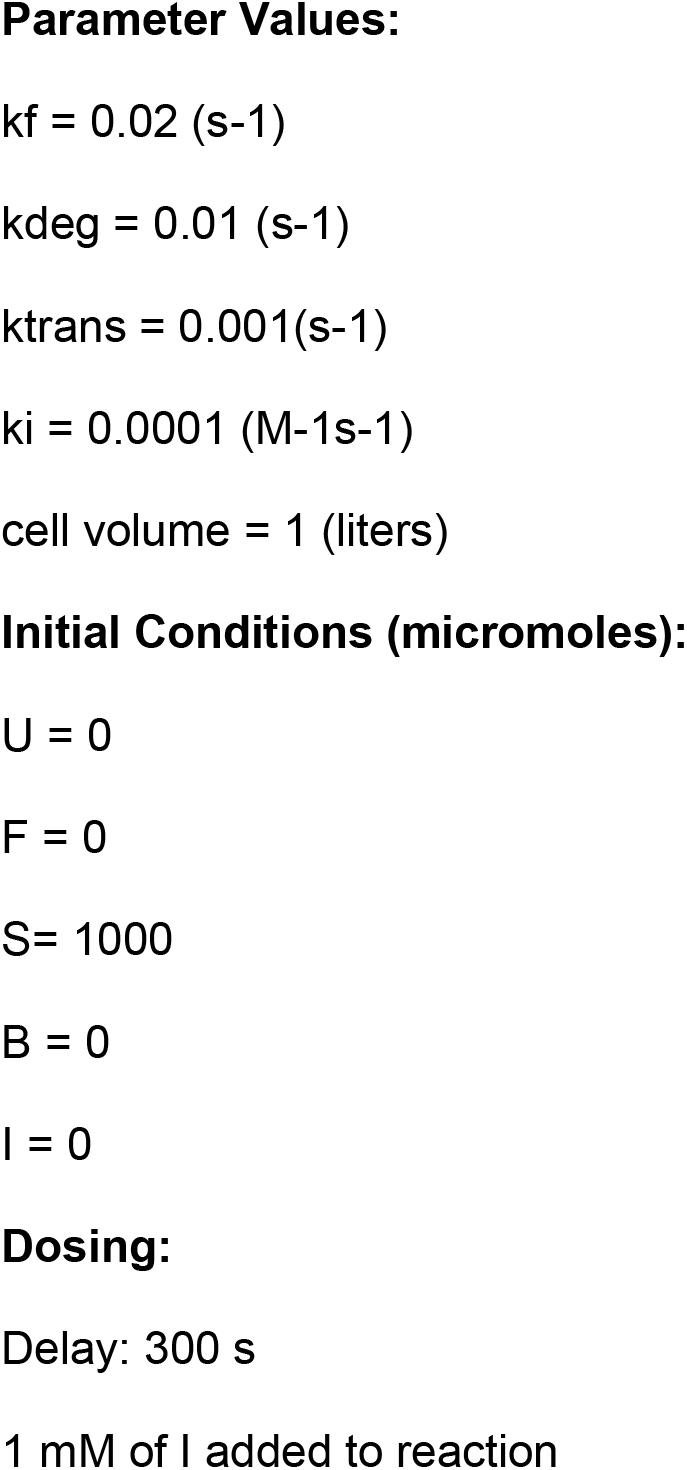
ODEs used for the kinetic simulations. (Related to figure 4) Numerical simulation was set up with a fixed concentration of mRNA/ribosome complex (S). These complexes could either form nascent polypeptides (U) with a rate ktrans that is a collective rate constant for transcription/translation, or be inhibited with a rate constant kblock in the presence of a translation inhibitor (I). The pool of U could either degrade with a rate kdeg or fold with a rate constant of kf and reach the native state (F). We finally monitor the total amount of undegraded protein (U+F) after blocking translation after 300 s of starting the simulation (simulation start mimics induction). The reaction volume was kept at 1 liter.

**Table S3:** Concentration of metabolites added to LB to measure effect of metabolites on Fluorescence *in vivo* (Related to Figure 7)

| S. no. | Metabolite | Concentration (mM) |
| --- | --- | --- |
| 1 | Alanine | 200 |
| 2 | Arginine | 2.5 |
| 3 | Betaine | 200 |
| 4 | Proline | 200 |
| 5 | Serine | 200 |
| 6 | TMAO | 200 |
| 7 | Aspartate | 200 |
| 8 | Malate | 100 |
| 9 | Malonate | 100 |
| 10 | Fumarate | 100 |
| 11 | Succinate | 100 |
| 12 | KCL | 350 |
| 13 | Sucrose | 400 |
| 14 | Sorbitol | 400 |

**Table S4:** Identity of genes along with the type of mutation observed in the evolved thermotolerant strains E5 and H8 on genome sequencing (Related to Figure 8)

| **Strain** | **Gene** | **Type** |
| --- | --- | --- |
| H8 | mngA | Ins del |
| H8 | hyfB | Ins del |
| H8 | fucl | Ins del |
| H8 | kefC | missense |
| H8 | sgrR | missense |
| H8 | mraY | missense |
| H8 | ftsW | missense |
| H8 | dxs | missense |
| H8 | cstA | stop |
| H8 | mngA | missense |
| H8 | clpA | missense |
| H8 | flgL | missense |
| H8 | oppB | missense |
| H8 | uidR | missense |
| H8 | otsA | missense |
| E5 | mngA | Ins del |
| E5 | csiR | Ins del |
| E5 | fucl | Ins del |
| E5 | rihC | missense |
| E5 | lptD | missense |
| E5 | mraY | missense |
| E5 | malZ | stop |
| E5 | cyoA | missense |
| E5 | paaH | missense |
| E5 | sdaA | missense |

